# The Dopamine Receptor Antagonist TFP Prevents Phenotype Conversion and Improves Survival in Mouse Models of Glioblastoma

**DOI:** 10.1101/870394

**Authors:** Kruttika Bhat, Mohammad Saki, Erina Vlashi, Fei Cheng, Sara Duhachek-Muggy, Claudia Alli, Garrett Yu, Paul Medina, Ling He, Robert Damoiseaux, Matteo Pellegrini, Nathan R. Zemke, Phioanh Leia Nghiemphu, Timothy F. Cloughesy, Linda M. Liau, Harley I. Kornblum, Frank Pajonk

## Abstract

Glioblastoma (GBM) is the deadliest adult brain cancer and all patients ultimately succumb to the disease. Radiation therapy (RT) provides survival benefit of 6 months over surgery alone but these results have not improved in decades. We report that radiation induces a glioma-initiating cell phenotype and we have identified trifluoperazine (TFP) as a compound that interferes with this phenotype conversion. TFP caused loss of radiation-induced Nanog mRNA expression, activation of GSK3 with consecutive post-translational reduction in p-Akt, Sox2 and β-catenin protein levels. TFP did not alter the intrinsic radiation sensitivity of glioma-initiating cells (GICs). Continuous treatment with TFP and a single dose of radiation reduced the number of GICs *in vivo* and prolonged survival in syngeneic and patient-derived orthotopic xenograft (PDOX) mouse models of GBM. Our findings suggest that combination of a dopamine receptor antagonist with radiation enhances the efficacy of RT in GBM by preventing radiation-induced phenotype conversion of radiosensitive non-GICs into treatment resistant, induced GICs.

**Significance:** GBM is the most common and most deadly adult brain cancer. The current standard-of-care is surgery followed by RT and temozolomide, which results in a median survival time of only 15 months. The efficacy of chemotherapies and targeted therapies in GBM is very limited because most of these drugs do not pass the blood-brain-barrier. Ultimately, all patients succumb to the disease. Our study describes radiation-induced cellular plasticity as a novel resistance mechanism in GBM. We identified a dopamine receptor antagonist as a readily available, FDA-approved drug known to penetrate the blood-brain-barrier which prevents phenotype conversion of glioma cells into glioma-initiating cells and prologs survival in mouse models of GBM, thus suggesting that it will improve the efficacy of RT without increasing toxicity.

## Introduction

GBM is among the deadliest cancer in adults with almost all patients succumbing to the disease. The median survival is only 15 months (1). The current standard-of-care for GBM patients involves postoperative RT (RT), which prolongs survival by ∼ 6 months over surgery alone (2, 3), and temozolomide treatment, which prolongs progression-free survival (PFS) by another ∼3 months over postoperative RT alone (1). Addition of classical chemotherapies, anti-angiogenic strategies and novel biologics to the current standard-of-care have all failed to improve the outcome for GBM patients, which has remained largely unchanged since the 1970s (2, 3).

Over the last four decades RT for GBM underwent several iterations from whole brain irradiation (WBI) versus gross tumor volume, standard fractionation schemes versus accelerated hyper-fractionated RT, and hypofractionated intensity-modulated radiation therapy (IMRT) (4). However, although RT is an indispensable part of the treatment regimen for GBM patients, strategies aimed at improving response to RT have failed to increase progression-free survival. The current standard-of-care in the RT of GBM has its origin in studies showing that escalating the WBI dose from 45 to 60 Gy led to increased survival (3). Yet, dose escalation from 60 Gy to 70, 80 and 90 Gy given to the tumor alone showed no dose response effect or improved outcome, and the tumors still recurred locally (5).

The almost universal fatal outcome of GBM patients treated by standard-of-care including postoperative RT stands in contrast to the intrinsic radiation sensitivity of GBM cells, which falls into the same range of radiosensitivity seen in other solid tumors frequently cured by total radiation doses of 60 Gy (6). In the above study, the SF_2Gy_ (surviving fraction at 2 Gy) values for glioma cells *in vitro* were comparable to those of other cancers including squamous cell carcinoma, colon cancer, and soft tissue sarcoma (6). Likewise, TCD_50_ (the radiation dose necessary to control 50% of the tumors locally) values were not exceptionally high and TCD_50_ and SF_2Gy_ values did not correlate with each other (6). With most systemic agents being ineffective and surgery and RT having reached their maximum potential, treatment for GBM has hit a critical barrier.

A mounting body of evidence points to a hierarchical organization of GBM with a small number of GICs at the apex of that hierarchy, able to self-renew and to repopulate a tumor (7, 8). Importantly, GICs were found to be relatively resistant to radiation (9) and chemotherapy (10). A number of marker systems have been used to prospectively identify GICs including the surface marker CD133 (9) and the lack of proteasome activity (11). A competing stochastic model assumes that every cell in GBM can acquire GIC traits over time. More recent reports indicate that while the hierarchical model describes the organization of GBM quite accurately, glioma cells exhibit plasticity that allows them to phenotype convert into induced GICs in response to stress (12).

In this study we identify radiation-induced phenotype conversion of GBM cells into radioresistant induced glioma-initiating cells as a novel facet of GBM treatment resistance. Importantly, we demonstrate that TFP, a compound identified in a high-throughput screen for its ability to prevent radiation-induced phenotype conversion of glioma cells into radiation-resistant tumor-initiating cells, can prolong survival in mouse models of GBM.

## Material and Methods

### Cell culture

Primary human glioma cell lines were established at UCLA as described (Hemmati et al., PNAS 2003 (7); Characteristics of specific gliomasphere lines can be found in Laks et al., Neuro-Oncology 2016 (13)). The GL261 murine glioma cell line was a kind gift of Dr. William H. McBride (Department of Radiation Oncology at UCLA). GL261 cells were cultured in log-growth phase in DMEM (Invitrogen, Carlsbad, CA) supplemented with 10% fetal bovine serum, penicillin and streptomycin. Primary GBM cells were propagated as gliomaspheres in serum-free conditions in ultra-low adhesion plates in DMEM/F12, supplemented with B27, EGF, bFGF and heparin as described previously (7, 11, 13). All cells were grown in a humidified atmosphere at 37°C with 5% CO_2_. The unique identity of all patient-derived specimens was confirmed by DNA fingerprinting (Laragen, Culver City, CA). All lines were routinely tested for mycoplasma infection (MycoAlert, Lonza).

ZsGreen-cODC expressing cells were obtained as described in (11). Briefly, cells were infected with a lentiviral vector coding for a fusion protein between the fluorescent protein ZsGreen and the C-terminal degron of murine ornithine decarboxylase (cODC). The latter, targets ZsGreen to ubiquitin-independent degradation by the 26S proteasome, thus reporting lack of proteasome function through accumulation of ZsGreen-cODC. We previously reported that cancer cell populations lacking proteasome activity are enriched for tumor-initiating cells in GBM, breast cancer and cancer of the head and neck region (11, 14, 15), and others have confirmed these findings independently in tumors of the liver, lung, cervix, pancreas, osteosarcoma and colon (16–21). After infection with the lentivirus, cells expressing the ZsGreen-cODC fusion protein were further selected with G418 for 5 days. Successful infection was verified using the proteasome inhibitor MG132 (Sigma, MO).

### Cell cycle analysis

After 5 days of 0 or 8 Gy irradiation, HK-374 ZsGreen-cODC expressing cells were stained with Hoechst 33342 and pyronin Y. Briefly, cells were trypsinized and rinsed with HBSS/1 mM HEPES/ 10% FBS. DNA was stained with 1 μg/ml Hoechst 33342 in HBSS/ 1 mM HEPES/10% FBS solution/50 μM Verapamil for 45 minutes at 37°C. The cells were rinsed once and then stained for RNA with 4 μM pyronin Y in HBSS/ 1 mM HEPES/10% FBS solution for 45 minutes at 37°C. Finally, cells were rinsed and re-suspended in PBS. At least 100,000 cells were analyzed by flow cytometry.

### Senescence analysis

HK-374 ZsGreen-cODC expressing cells were plated in 6 well plates at a density of 50,000 cells/well and the next day irradiated with 0 or 8 Gy. 5 days after irradiation the plates were assayed for senescence via the Senescence β-galactosidase Cell Staining Kit (Cell Signaling, Catalog # 9860) and the staining was performed as per manufacturer’s protocol.

### Animals

6–8-week-old C57BL/6 mice, or NOD-*scid* IL2Rgamma^null^ (NSG) originally obtained from The Jackson Laboratories (Bar Harbor, ME) were re-derived, bred and maintained in a pathogen-free environment in the American Association of Laboratory Animal Care-accredited Animal Facilities of Department of Radiation Oncology, University of California (Los Angeles, CA) in accordance with all local and national guidelines for the care of animals. Weights of the animals were recorded every day. 2×10^5^ GL261-Luc and 3×10^5^ HK-308-Luc or HK-374-Luc cells were implanted into the right striatum of the brains of mice using a stereotactic frame (Kopf Instruments, Tujunga, CA) and a nano-injector pump (Stoelting, Wood Dale, IL). Injection coordinates were 0.5 mm anterior and 2.25 mm lateral to the bregma, at a depth of 3.5 mm from the surface of the brain. Tumors were grown for 3 (HK-374), 7 (GL261) or 21 (HK-308) days after which successful grafting was confirmed by bioluminescence imaging. Mice that developed neurological deficits requiring euthanasia were sacrificed.

### In vivo bioluminescent imaging

Starting one and three weeks after implantation of xenografts, GL261-Luc-bearing C57BL/6 mice and NSG mice bearing HK-308-Luc or HK-374-Luc tumors were imaged at regular intervals and the tumor-associated bioluminescent signal was recorded. Prior to imaging the mice were injected intra-peritoneally with 100µl of D-luciferin (15 mg/ml, Gold Biotechnology). Five minutes later, animals were anesthetized (2 % isofluorane gas in O_2_) and luminescence was recorded (IVIS Spectrum, Perkin Elmer, Waltham, MA). Images were analyzed with Living Image Software (Caliper LifeSciences).

### Brain tissue digestion and flow cytometry

2×10^5^ GL261-StrawberryRed and 3×10^5^ HK-374- or HK-157-StrawberryRed cells were implanted into the brains of C57BL/6 or NSG mice respectively, as described above. Tumors were grown for 3 (HK-374) and 7 days (HK-157 and GL261) for successful grafting. Mice-bearing tumors were injected intra-peritoneally on a 5-days on / 2-days off schedule for 1 or 2 weeks (GL261), 4 weeks (HK-374) and 6 weeks (HK-157) either with TFP or saline. TFP was dissolved in sterile saline at a concentration of 2.5 mg/mL. All animals were treated with 20 mg/kg TFP. At the indicated time points after implantation the mice were sacrificed and tumor-bearing brains were dissected for further analysis. Detailed procedure for brain tumor dissociation and flow cytometric analysis is available in *SI Appendix*.

For radiation-induced reprogramming experiments *in vivo* with GL261-StrawberryRed and GL261-BFP cells, GL261-StrawberryRed cells were first stained with an anti-Prominin-APC antibody (Miltenyi, Cat # 130-102-197) and the Prominin-positive cells were removed by FACS. The Prominin-negative cells were seeded into 6-well plates at a density of 50,000 cells/well. Cells were irradiated with 0 or 4 Gy and incubated at 37°C in a CO_2_ incubator for five days. After 5 days, GL261-StrawberryRed cells along with GL261-BFP cells were stained with an anti-Prominin-APC antibody to determine the percentage of prominin-positive cells in the total population. 25,000 cells from each cell population (GL261-StrawberryRed and GL261-BFP) and from each group (0 Gy or 4 Gy) were mixed together and a total of 50,000 cells were injected into mice intracranially. The mice were maintained for 3 weeks and after that the brains were dissociated as described above for flow cytometric analysis with GL261-StrawberryRed and GL261-BFP cells stained with anti-Prominin antibody.

### High-throughput screening

For primary screening of drug libraries ZsGreen-cODC-negative SUM159PT cells (Asterand Bioscience, Cambridge, MA), which show high levels of radiation-induced phenotype conversion (22) were used as a discovery platform. Cells were sorted by high-speed FACS (FACS Aria I/II) in the UCLA Jonsson Comprehensive Cancer Center (JCCC) and Center for AIDS Research Flow Cytometry Core Facility.

Concurrently, low evaporation 384-well plates were filled by a manifold liquid dispenser (Multidrop 384, Thermo LabSystems) with 35µL media/well (F-12, hydrocortisone (100mg/2mL), 5% FBS, 1 % penicillin-streptomycin, 0.1% insulin, and 1M HEPES). Using a liquid handler (Biomek FX, Beckman Coulter, Brea, CA) with custom pin tool (V&P scientific, San Diego, CA), 320 unique compounds (500 nL each from 1mM stocks in neat DMSO) were then transferred to each 384-well plate according to pre-defined plate maps. The other 64 wells received an equal volume of DMSO alone, serving as negative controls. Following re-suspension of cells in media at a concentration of 200,000 cells/mL, 15µL of cell suspension (3,000 cells) was plated into each well of the pre-filled 384-well plates. After completion of cell plating, the 384-well plates were kept at 37°C, 5% CO_2_.

Twenty-four hours after plating, cells were irradiated with 8Gy. Five days later, 10 µL Hoechst 33342 solution (25 µg/mL) was added to the cells. The plates were incubated at 37°C, 5% CO_2_ for one hour. Plates were scanned using an Acumen Mark III laser-scanning cytometer cytometer (TTP Labtech, Melbourn, UK). Laser scanning with a 488 nm laser was used for detection of cells expressing the fusion protein, ZsGreen-cODC. Additionally, scanning with a UC laser at 405 nm allowed for detection of Hoechst-stained nuclei, giving a measure of total viable cells. Calculation for z-score statistics is available in *SI Appendix*.

### Drug treatment

After confirming tumor grafting via bioluminescent imaging, mice bearing GL261 tumors were injected intra-peritoneally on a 5-days on / 2-days off schedule for 3 weeks either with TFP or saline. Mice implanted with the HK-374 or HK-308 specimen were injected subcutaneously on a 5-days on / 2-days off schedule with TFP or saline until they reached euthanasia endpoints. TFP was dissolved in sterile saline at a concentration of 2.5 mg/mL. All animals were treated with 20 mg/kg TFP, the published MTD for mice (23).

### Irradiation

Cells or mice were irradiated at room temperature using an experimental X-ray irradiator (Gulmay Medical Inc. Atlanta, GA) at a dose rate of 5.519 Gy/min for the time required to apply a prescribed dose. The X-ray beam was operated at 300 kV and hardened using a 4 mm Be, a 3 mm Al, and a 1.5 mm Cu filter and calibrated using NIST-traceable dosimetry. Corresponding controls were sham irradiated.

For the assessment of the effect of TFP in combination with irradiation *in vivo*, mice were anesthetized prior to irradiation with an intra-peritoneal injection of 30 uL of a ketamine (100 mg/mL, Phoenix, MO) and xylazine (20 mg/mL, AnaSed, IL) mixture (4:1) and placed on their sides into an irradiation jig that allows for irradiation of the midbrain while shielding the esophagus, eyes and the rest of the body. Animals implanted with GL261 cells received a single dose of 10 Gy on day 8 after tumor implantation. Animals injected with the HK-308 glioma specimen received a single dose of 4 or 10 Gy on day 21 after tumor implantation. Animals injected with the HK-374 glioma specimen received a single dose of 10 Gy on day 3 after tumor implantation.

### *In-vitro* sphere formation assay

For the assessment of self-renewal *in vitro*, cells were irradiated with 0, 2, 4, 6 or 8 Gy and seeded under serum-free conditions into plates, not treated for tissue culture, in DMEM/F12 media, supplemented with 10 mL / 500 mL of B27 (Invitrogen), 0.145 U/mL recombinant insulin (Eli Lilly, Indiana), 0.68 U/mL heparin (Fresenius Kabi, Illinois), 20 ng/mL fibroblast growth factor 2 (bFGF, Sigma) and 20 ng/mL epidermal growth factor (EGF, Sigma). The number of spheres formed at each dose point was normalized against the non-irradiated control. The resulting data points were fitted using a linear-quadratic model.

### Quantitative Reverse Transcription-PCR

Total RNA was isolated using TRIZOL Reagent (Invitrogen). cDNA synthesis was carried out using the SuperScript Reverse Transcription III (Invitrogen). Quantitative PCR was performed in the My iQ thermal cycler (Bio-Rad, Hercules, CA) using the 2x iQ SYBR Green Supermix (Bio-Rad). *C*_t_ for each gene was determined after normalization to GAPDH or RPLP0 and ΔΔ*C*_t_ was calculated relative to the designated reference sample. Gene expression values were then set equal to 2^−ΔΔCt^ as described by the manufacturer of the kit (Applied Biosystems). All PCR primers were synthesized by Invitrogen. Primer list and the primer sequences are available in *SI Appendix*.

### Chromatin Immunoprecipitation Polymerase Chain Reaction (ChIP-PCR)

ChIP-PCR was performed using SimpleChIP^®^ Plus Enzymatic Chromatin IP Kit (Cell Signaling, Cat # 9005) by following the manufacturer’s protocol. Briefly, HK-374 cells were plated in 15 cm dishes and cultured until they reached confluency. The plates were irradiated at 0 or 4 Gy and 48 hours later these cells were used to perform the ChIP-PCR. Detailed description of the ChIP-PCR is available in *SI Appendix*.

### Histone Enzyme-Linked Immunosorbent Assay

HK-374 cells were irradiated with 0 and 4 Gy. Fourty-eight hours after irradiation histone proteins were isolated using EpiQuik Total Histone Extraxtion Kit (Epigentek, Farmingdale, NY, Cat #OP-0006). Detailed description of the method is available in *SI Appendix*.

### RNASeq

One and 48 hours after 4 Gy irradiation or sham irradiation, RNA was extracted from ZsGreen-cODC-negative HK-374 non-GICs using Trizol. RNASeq analysis was performed by Novogene (Chula Vista, CA). Quality and integrity of total RNA was controlled on an Agilent Technologies 2100 Bioanalyzer (Agilent Technologies; Waldbronn, Germany). The RNA sequencing library was generated using NEBNext® Ultra RNA Library Prep Kit (New England Biolabs) according to manufacturer’s protocols. The library concentration was quantified using a Qubit 3.0 fluorometer (Life Technologies), and then diluted to 1 ng/uL before checking insert size on an Agilent Technologies 2100 Bioanalyzer (Agilent Technologies; Waldbronn, Germany) and quantifying to greater accuracy by quantitative Q-PCR (library molarity >2 nM). The library was sequenced on Illumina NovaSeq6000 with an average of 43.4M reads per RNA sample. Detailed description on downstream analysis and differential expression analysis is available in *SI Appendix*.

### Assay for Transposase-Accessible Chromatin using sequencing (ATAC-Seq)

Cells were harvested and frozen in culture media containing fetal bovine serum (FBS) and 5% DMSO. Cryopreserved cells were sent to Active Motif to perform the ATAC-seq assay. The cells were then thawed in a 37°C water bath, pelleted, washed with cold PBS, and tagmented as previously described (24), with some modifications based on (25). Briefly, cell pellets were resuspended in lysis buffer, pelleted, and tagmented using the enzyme and buffer provided in the Nextera Library Prep Kit (Illumina). Tagmented DNA was then purified using the MinElute PCR purification kit (Qiagen), amplified with 10 cycles of PCR, and purified using Agencourt AMPure SPRI beads (Beckman Coulter). Resulting material was quantified using the KAPA Library Quantification Kit for Illumina platforms (KAPA Biosystems), and sequenced with PE42 sequencing on the NextSeq 500 sequencer (Illumina).

Analysis of ATAC-seq data was very similar to the analysis of ChIP-Seq data. Reads were aligned to the human genome (hg38) using the BWA algorithm (mem mode; default settings). Duplicate reads were removed, only reads mapping as matched pairs and only uniquely mapped reads (mapping quality ≥ 1) were used for further analysis. Alignments were extended *in silico* at their 3’-ends to a length of 200 bp and assigned to 32-nt bins along the genome. The resulting histograms (genomic “signal maps”) were stored in bigWig files. Peaks were identified using the MACS 2.1.0 algorithm at a cutoff of p-value 1e-7, without control file, and with the –nomodel option. Peaks that were on the ENCODE blacklist of known false ChIP-Seq peaks were removed. Signal maps and peak locations were used as input data to Active Motifs proprietary analysis program, which creates Excel tables containing detailed information on sample comparison, peak metrics, peak locations and gene annotations.

### siRNA Treatment

70% confluent HK-374 cell cultures were grown in antibiotic-free culture media overnight. The next day the culture media was replaced with 1 ml of Opti-MEM™ reduced serum media. To this control or DRD2 siRNA-lipid complex (1:1) was added as per the manufacturer’s protocol. Detailed description of siRNA treatment and protein knockdown validation is available in *SI Appendix*.

### Western Blotting

GBM cells were serum starved overnight and the following day pre-treated with 10 μM TFP for one hour. Pre-treated HK-374 and HK-345 cells were irradiated with a single dose of 8 Gy immediately after a second treatment with 10 μM TFP. One hour after irradiation, the cells were lysed in RIPA lysis buffer containing proteinase inhibitor and phosphatase inhibitor. The protein concentration in each sample was determined by BCA protein assay and samples were denaturated in 4X Laemmli sample buffer containing 10% β-mercaptoethanol for 10 minutes at 95°C. Western blotting was performed to demonstrate the change in protein expression levels from different treatment conditions.

To analyze for changes in the expression levels of Yamanaka factors in HK-374 cells five days after exposing them to 8 Gy irradiation, Western blotting was performed using proteins extracted from FACS sorted ZsGreen-cODC-negative and ZsGreen-cODC-positive cells treated with 8 Gy. Detailed description of the Western blotting procedure and the protein targets is available in *SI Appendix*.

### Immunohistochemistry

Brains were explanted, fixed in formalin for twenty-four hours and embedded in paraffin. Immunohistochemistry was performed on these paraffin-embedded slides. The method used is available in *SI Appendix*.

### Statistics

Unless stated otherwise all data shown are represented as mean ± standard error mean (SEM) of at least 3 biologically independent experiments. A *p*-value of ≤0.05 in an unpaired two-sided *t*-test indicated a statistically significant difference. Kaplan-Meier estimates were calculated using the GraphPad Prism Software package. For Kaplan-Meier estimates a *p*-value of 0.05 in a log-rank test indicated a statistically significant difference.

### Data sharing

All data discussed in this manuscript will be made available upon reasonable request. The RNA-Seq data is available via GEO accession number GSE166358.

## Results

### Relative radiation sensitivity of glioma-initiating cells

As described before, the radiation sensitivity of bulk glioma cell populations was found to be in the same range as the radiosensitivity of other solid tumors frequently cured by radiation (6, 26). In light of the known heterogeneity of GBM cell populations and the reported relative radioresistance of glioma-initiating cells compared to their more differentiated progeny (9), we first sought to test if the radioresistance of GICs differed from that of tumor-initiating cells in breast cancer. Using 4 primary patient-derived GBM lines we performed sphere formation assays under serum-free conditions after exposing the cells to radiation doses of 0, 2, 4, 6 or 8 Gy (**Figure 1A**). Under these conditions, GICs form gliomaspheres while more differentiated glioma cells die from anoikis. As expected, GICs exhibited a range of surviving fractions of 0.4 to 0.63 at 2 Gy. However, a comparison with our published data on the radiation sensitivity of breast cancer-initiating cells (27) did not reveal a more radioresistant phenotype of GICs. Like the radiation sensitivity of bulk cell populations in GBM which falls into the same range of radiation sensitivity reported for other solid cancers (6), the radiation sensitivity of GICs is comparable to that of their equivalent in breast cancer, thus suggesting that the intrinsic radiation resistance of GICs does not explain why GBM cannot be controlled by RT and often recurs within or in proximity to the radiation therapy target volume.

**Figure 1.**
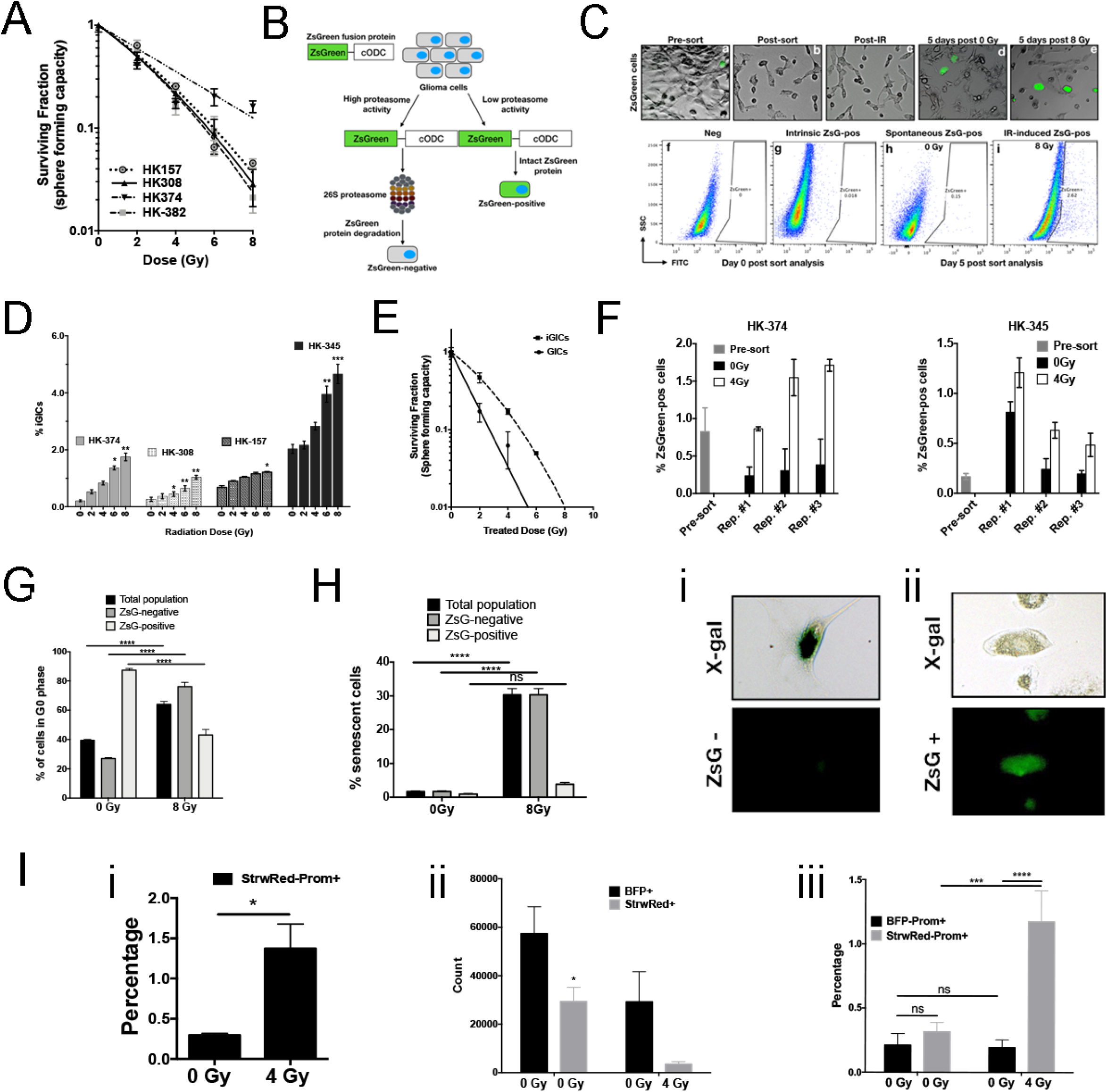
Radiation induces phenotype conversion in GBM. **(A)** Four primary patient-derived GBM lines (parent HK-157, HK-308, HK-374 and HK-382) were seeded as single cell suspension cultures in 96-well plates containing serum-free medium and exposed to radiation doses of 0, 2, 4, 6 or 8 Gy. The spheres were maintained by feeding with fresh growth factors (10X medium) regularly. The number of spheres formed at each dose point was counted and normalized against their respective non-irradiated control. The final curve was generated using a linear-quadratic model. **(B)** Schematical representation of the ZsGreen-cODC reporter system. **(C)** Representative images (10X) of HK-374 ZsGreen-cODC infected cells before and after sort (a & b), after irradiation (c), 5 days after 0 Gy irradiation (d) and 5 days post 8 Gy irradiation (e). Flow sort analysis representative images of HK-374-negative cells (f), HK-374 cells before sorting (g), 0 Gy sample 5 days post irradiation (g) and 8 Gy samples 5 days post sorting (h). **(D)** Four primary patient-derived GBM lines (HK-308, HK-374, HK-157 and HK-345) were infected with ZsGreen-cODC reporter vector and maintained as an adherent culture in a 10% serum containing medium. After selection, the cells were sorted to deplete the ZsGreen-cODC-positive GICs. The remaining population consisting of differentiated ZsGreen-cODC-negative glioma cells were plated at a density of 50,000 cells per well in a 6-well plate and the following day irradiated with single doses of 0, 2, 4, 6 or 8 Gy. Five days later, the cells were detached and analyzed for ZsGreen-cODC-positive cells by flow cytometry, using their non-infected parent lines as controls, and represented as percentage iGICs. **(E)** ZsGreen-cODC vector expressing HK-374 cells were sorted and the ZsGreen-cODC-positive cells obtained were seeded in a 96-well plate for sphere formation assay. The remaining population was plated and subjected to irradiation at 0, 2, 4, 6 or 8 Gy. Five days later, the cells were detached and sorted for ZsGreen-cODC positive cells. These iGICs were then seeded in a 96-well plate for sphere formation assay. The number of spheres formed at each dose point was counted and normalized against the control. The resulting data was fitted using the linear-quadratic model. **(F)** ZsGreen-cODC vector expressing HK-374 and HK-345 cells were sorted for ZsGreen-cODC-negative cells and irradiated at 4 Gy. Five days later, a small portion of these cells were used for flow cytometry to analyze for percentage of ZsGreen-cODC-positive cells. The remaining cells were sorted for ZsGreen-cODC-negative cells and again irradiated at 4 Gy. This approach was followed two more times. The ZsGreen-cODC positive cells obtained after each repeat was graphed and presented as percentage ZsGreen-cODC-positive cells. **(G)** Cell cycle analysis demonstrating percentage of cells in G_0_ phase of the cell cycle in three different cell populations - total, ZsGreen-cODC-negative and ZsGreen-cODC-positive cells, after 5 days of irradiation with a single dose of 0 and 8 Gy irradiation. **(H)** Cellular senescence analysis five days post single dose of 0 and 8 Gy irradiation in HK-374 ZsGreen-cODC infected cells demonstrating percentage of senescent cells in three different populations - total, ZsGreen-negative and ZsGreen-cODC-positive cells. Representative images of a cell showing positive for X-gal but negative for ZsGreen-cODC protein (i); representative image of a cell showing negative for X-gal but positive for ZsGreen-cODC protein (ii). **(I)** GL261-StrawberryRed cells were first sorted for cells negative for Prominin. The sorted cells were then irradiated with a single dose of 4 Gy and incubated at 37°C in a CO_2_ incubator for 5 days. Percentage of Prominin^+^ cells in the cell preparation from the pre-sort (0 Gy) and the sorted, irradiated and 5 days post incubation (4 Gy) of GL261-sorted-irradiated-StrawberryRed cells were analyzed by performing flow cytometry (i). Next, 1:1 ratio of the sorted GL261-StrawberryRed cells and the unsorted GL261-BFP cells were intracranially implanted into C57Bl/6 mice. Three weeks post implantation, the brains of the mice were extracted, tumor cells were dissociated, and labeled with anti-promonin antibody. The antibody labeled cells were subjected to flow cytometric analysis and the number of either StrawberryRed or BFP-positive cells obtained from each brain dissociation prep was graphed (ii); the cells positive for both StrawberryRed and Prominin or BFP and Prominin were also graphed and presented as percentage positive (iii). All experiments in this figure have been performed with at least 3 biological independent repeats. *p*-values were calculated using unpaired t-test. * *p*-value < 0.05, ** *p*-value < 0.01, *** *p*-value < 0.001 and **** *p*-value < 0.0001.

### Radiation-induced phenotype conversion in GBM

We have previously reported that triple-negative and claudin-low breast cancers exhibit high rates of spontaneous and radiation-induced phenotype conversion of non-tumorigenic breast cancer cells into breast-cancer initiating cells, while non-tumorigenic luminal breast cancer cells, show only very low rates of phenotype conversion (22). Likewise, phenotype conversion from CD133^neg^ into CD133^pos^ GICs in response to changes in the tumor microenvironment has been previously reported for GBM (12). Phenotype conversion in irradiated glioma cells has not been investigated and therefore, we next tested if ionizing radiation would also induce this phenomenon in GBM. We utilized our imaging system for tumor-initiating cells (11) to deplete cell populations from ZsGreen-cODC-positive GICs with low proteasome activity by high-speed FACS (**Figure 1B**). The system is based on a fusion protein of ZsGreen and the C-terminal degron of murine ornithine decarboxylase (cODC), which targets the protein to ubiquitin-independent proteasomal degradation. Cell populations lacking proteasome activity can be identified by accumulation of the fluorescent ZsGreen protein and are enriched for tumor-initiating cells in a large number of solid tumors including GBM (11, 15, 28). In order to confirm our prior results, we implanted ZsGreen-positive cells into immunodeficient mice and found that they formed more aggressive tumors *in vivo* (**Supplementary Figure 1**). Once GICs were removed by sorting, the remaining populations of differentiated glioma cells were irradiated with single doses of 0, 2, 4, 6, or 8 Gy. Five days after a single dose of radiation, the cells were analyzed by flow cytometry and the emergence of ZsGreen-cODC-positive cells with low proteasome activity was interpreted as a measure for phenotype conversion. In general, the patient-derived primary GBM samples displayed various degrees of spontaneous (without irradiation treatment) phenotype conversion in the range of 0.21 to 2 % (**Figure 1C/D**, 0 Gy). Irradiation with single doses led to a dose-dependent increase of phenotype conversion up to 4.7 % of the cells undergoing phenotype conversion after 8 Gy (**Figure 1C/D**). Importantly, induced GICs were more resistant to radiation than preexisting GICs *in vitro* (**Figure 1E**).

Next, we tested if phenotype conversion could be repeatedly induced or if it rather reflected a population of cells prone to acquiring GIC traits. ZsGreen-cODC-positive cells were purged from the bulk population of HK-374 and HK-345 cells by FACS and the remaining populations were irradiated with 4 Gy. After five days, the number of radiation-induced ZsGreen-cODC-positive (iGICs) was assessed, ZsGreen-cODC-positive cells were purged again and the remaining cells re-irradiated with 4 Gy and tested for the induction of ZsGreen-cODC-positive iGICs five days later. Using this approach, we found that radiation-induced phenotype conversion was not restricted to a single initial irradiation but could be repeatedly observed, thus indicating that the process cannot simply be explained by cell selection or reflect a restriction to a preexisting cell population prone to phenotype conversion (**Figure 1F**).

Furthermore, we addressed if radiation would increase the number of quiescent ZsGreen-cODC positive cells by staining the RNA and DNA content of the cells. In agreement with the literature, radiation caused cell cycle arrest in the bulk population of ZsGreen-cODC-negative cells. Unirradiated ZsGreen-cODC-positive cells were predominantly in a quiescent state (G_0_) but were recruited into the cell cycle in response to radiation (**Figure 1G**).

Treatment of unsorted bulk tumor cell population with radiation enriches for radioresistant tumor-initiating cells (9). In order to study the cell fate of ZsGreen-cODCpositive and –negative cells after irradiation, HK-374-ZsGreen-cODC cells were treated with 0 or 4 Gy. Five days later, cells were fixed and stained for beta-galactosidase activity, a marker for cellular senescence (29). Radiation significantly increased the number of senescent ZsGreen-cODC-negative cells but did not significantly increase the number of senescent ZsGreen-cODC-positive cells (**Figure 1H**). Together, this indicated that the radiation-induced occurrence of ZsGreen-cODC-positive cells from ZsGreen-cODC-negative cells cannot be explained by induction of senescence or quiescence.

To test if induced GICs contribute to the recurrence of GBM after irradiation, we engineered GL261 cells to constitutively express the fluorescent proteins Strawberry-Red (StrawRed) or Blue-fluorescent protein (BFP). Using CD133 as a validated marker for GICs in GL261 (30) we removed CD133-positive cells from StrawRed-expressing cells and irradiated the remaining CD133-negative cells with 0 or 4 Gy. After 5 days in culture, the percentage of CD133-positive cells was assessed by flow cytometry. Radiation induced a phenotype conversion of CD133-negative cells into CD133-positive cells that significantly exceeded the number of preexisting CD133-positive cells in unirradiated GL261 cells (**Figure 1I(i)**). 25,000 cells of irradiated StrawRed-expressing CD133-negative and 25,000 cells unirradiated, unsorted BFP-expressing cells were mixed and injected into the striatum of C57Bl/6 mice. After 3 weeks, the brains were harvested and analyzed for the number of CD133-positive StrawRed and BFP expressing cells. Irradiation with 4 Gy reduced the total number of grafting StrawRed-expressing cells, which was in line with the cytotoxic effects of radiation (**Figure 1I(ii)**). However, the population of grafting StrawRed cells was significantly enriched for CD133-positive induced GICs suggesting an *in vivo* contribution of iGICs to tumor recurrence after *in vitro* radiation (**Figure 1I(iii)**).

### Identification of compounds that interfere with phenotype conversion

With the exception of temozolomide, previous attempts to prolong survival of glioma patients through pharmacological intervention have largely failed. Clinically, the blood brain barrier limits the use of established chemotherapeutic agents in GBM, and GICs have been shown to resist most established systemic therapies (10). Cancer treatment-induced conversion of GBM cells into therapy-resistant GICs could potentially add to this resistance. Therefore, we sought to screen chemical libraries for compounds that interfere with this process. In a high-throughput screen of 83,000 compounds including libraries of FDA-approved drugs, we identified the dopamine receptor antagonist TFP as a potent inhibitor of radiation-induced phenotype conversion.

To confirm the presence of TFP’s target on GBM cells, we characterized expression levels of dopamine receptors (DRD1-4 relative to DRD5) in differentiated and matching GIC-enriched GBM cultures from the same GBM specimen via qRT-PCR. The paired specimen from the HK-308, HK-157 and HK-374 lines expressed at least one of the DRD receptors, while HK-345 cells had very low expression levels for all the DRD receptors relative to DRD5 (**Figure 2A**). Reflective of D_2_-like DRD expression status, TFP prevented radiation-induced phenotype conversion in HK-308, HK-157 and HK-374 specimens, but had no effect on HK-345 cells (**Figure 2B**) supporting a role for dopamine receptors in radiation-induced phenotype conversion in GBM.

**Figure 2.**
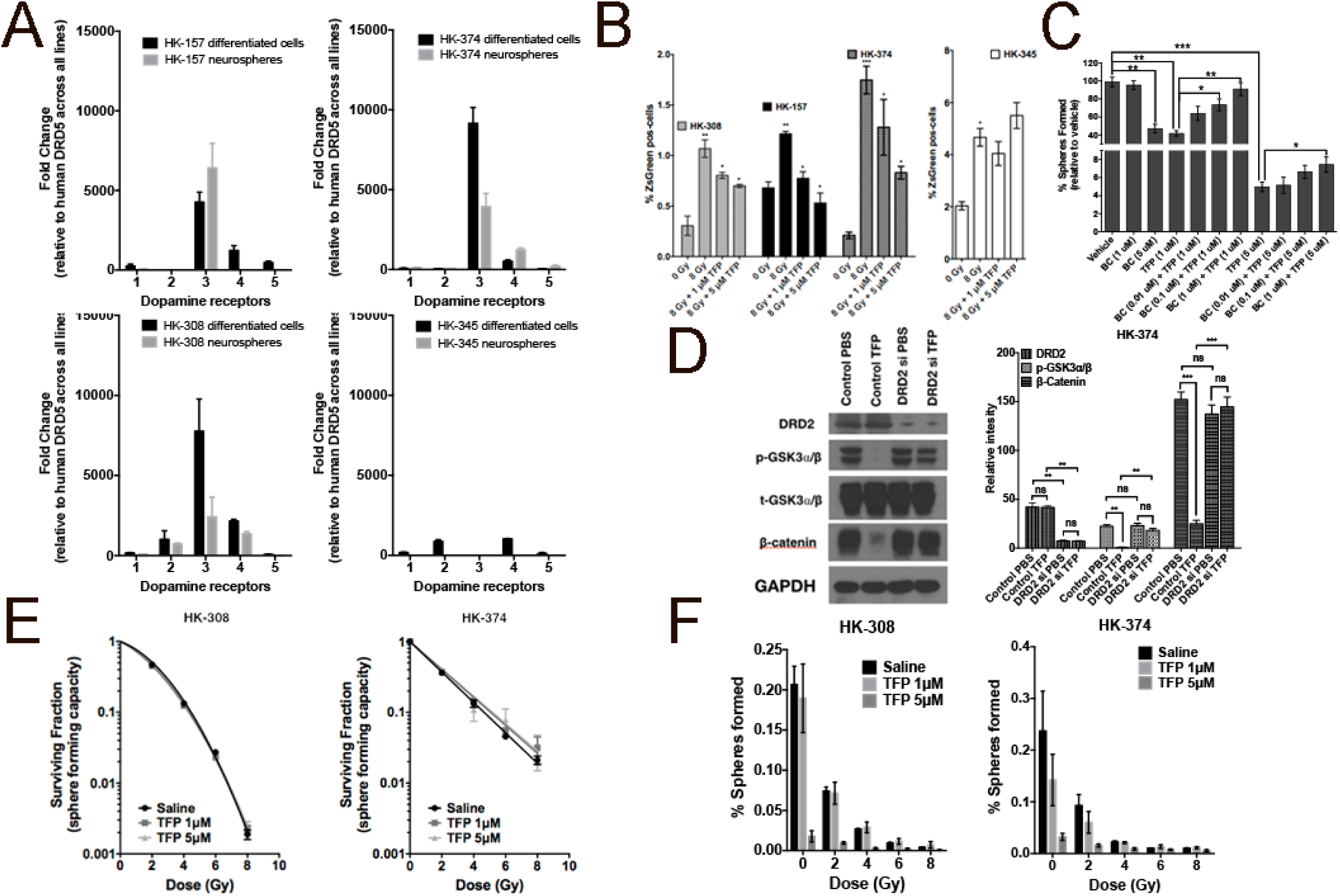
TFP prevents radiation-induced phenotype conversion in GBM. **(A)** Four primary patient-derived GBM lines (HK-157, HK-374, HK-308 and HK-345) were gown as differentiated cells in the presence of 10 % serum and as gliomaspheres in serum-free medium. RNA from both cultures was isolated and qPCR was performed to analyze for the difference in Dopamine Receptor (DRD1, DRD2, DRD3, DRD4 and DRD5) expression levels. GAPDH was used as the internal control to obtain the dCt values. The dCt values were normalized to DRD5 across all lines to obtain the ddCt values. The fold change in expression levels of DRDs was calculated by 2^-ddCt method. **(B)** Sorted ZsGreen-cODC-negative cells from HK-308, HK-157, HK-374 and HK-345 ZsGreen-cODC vector expressing cells were plated at a density of 50,000 cells per well in a 6-well plate and the following day pre-treated either with TFP (1 and 5 uM) or vehicle (Saline) one hour before irradiation at a single dose of 0 or 8 Gy. Five days later, the cells were detached and analyzed for ZsGreen-cODC-positive cells by flow cytometry, using their non-infected parent lines as controls, and represented as percentage iGICs. **(C)** Sphere formation assay was performed using the HK-308 parent gliomaspheres plated in a 96-well plate and treating them with different concentrations of TFP (1 and 5 uM) in combination with or without Bromocryptine (0.1, 1 and 5 uM). The spheres were fed with growth factors (10X) medium regularly. The number of spheres formed in each condition was counted and normalized against the vehicle control. The resulting data was presented as percentage spheres formed. **(D)** HK-374 cells were treated with either control siRNA or DRD2-specific siRNA for 72 hours, serum-starved for four hours and then treated with or without TFP (10 uM). Proteins were extracted and subjected to Western blotting. The membranes were blotted for p-GSK3a/b, t-GSK3a/b, β-catenin, and GAPDH. The intensity of each band was calibrated using ImageJ and presented as density ratio of gene over GAPDH in HK-374 cells. **(E & F)** Parent HK-308 and HK-374 gliomaspheres seeded in a 96-well plate were pre-treated either with TFP (1 or 5 uM) or vehicle (Saline) one hour before irradiation at a single dose of 0, 2, 4, 6 or 8 Gy. This set up was maintained with regular feeding of the spheres with 10X growth factors. The number of spheres formed in each condition was counted and normalized against the respective non-irradiated control. The final curve was generated using a linear-quadratic model. All experiments in this figure have been performed with at least 3 biological independent repeats. *p*-values were calculated using unpaired t-test. * *p*-value < 0.05, ** *p*-value < 0.01 and *** *p*-value < 0.001.

We further validated DRDs’ interfering with self-renewal by co-incubating cells with the DRD agonist, bromocriptine (BC). BC could reverse the inhibitory effect of TFP (**Figure 2C**). Next, we knocked down DRD2 expression in HK-374 cells. In wild-type cells, TFP caused a loss of Ser9/21 phosphorylation of GSK3α/β and subsequent degradation of β-catenin. Knock-down of DRD2 prevented TFP-induced loss of Ser9/21 phosphorylation and loss of β-catenin, thus demonstrating dependence of TFP’s effects on β-catenin on the presence of DRD2 (**Figure 2D**).

Using sphere-forming assays for HK-308 and HK-374 cells we further demonstrated that TFP did not act as a classical radiosensitizer as radiation survival curves of TFP-treated and irradiated samples were identical to their saline-treated corresponding controls (**Figure 2E**). However, when combined with radiation, TFP had an additive inhibitory effect on self-renewal capacity as was assessed in a sphere-formation assay (**Figure 2F**).

### TFP prevents radiation-induced re-expression of Yamanaka factors

We have previously shown that radiation-induced phenotype conversion in breast cancer coincided with the re-expression of the Yamanaka factors Sox2, Oct4, Klf4, c-Myc and their downstream target Nanog (22). Intrinsic GICs (ZsGreen-cODC-positive cells) showed elevated expression levels of Yamanaka factors when compared to ZsGreen-cODC-negative cells (**Figure 3A**, white bars). When patient-derived HK-374 GBM cells, depleted of ZsGreen-cODC-positive cells, were subjected to irradiation, the expression levels of Yamanaka factors, c-Myc, Oct4, Klf4 and Nanog were significantly elevated in the radiation-induced GICs over preexisting GICs (**Figure 3A**, checkered bars) and radiation-induced GICs showed elevated protein levels of Sox2, Oct4, Klf4 and Nanog when compared to GBM cells that did not convert their phenotype in response to radiation (**Fig. 3B**). Similarly, Yamanaka factor expression was upregulated in iGICs derived from HK-308 cells (**Figure 3C**, black bars). Importantly, TFP prevented the radiation-induced up-regulation of the Yamanaka factors and their downstream target, Nanog (**Figure 3C**, light gray bars).

**Figure 3.**
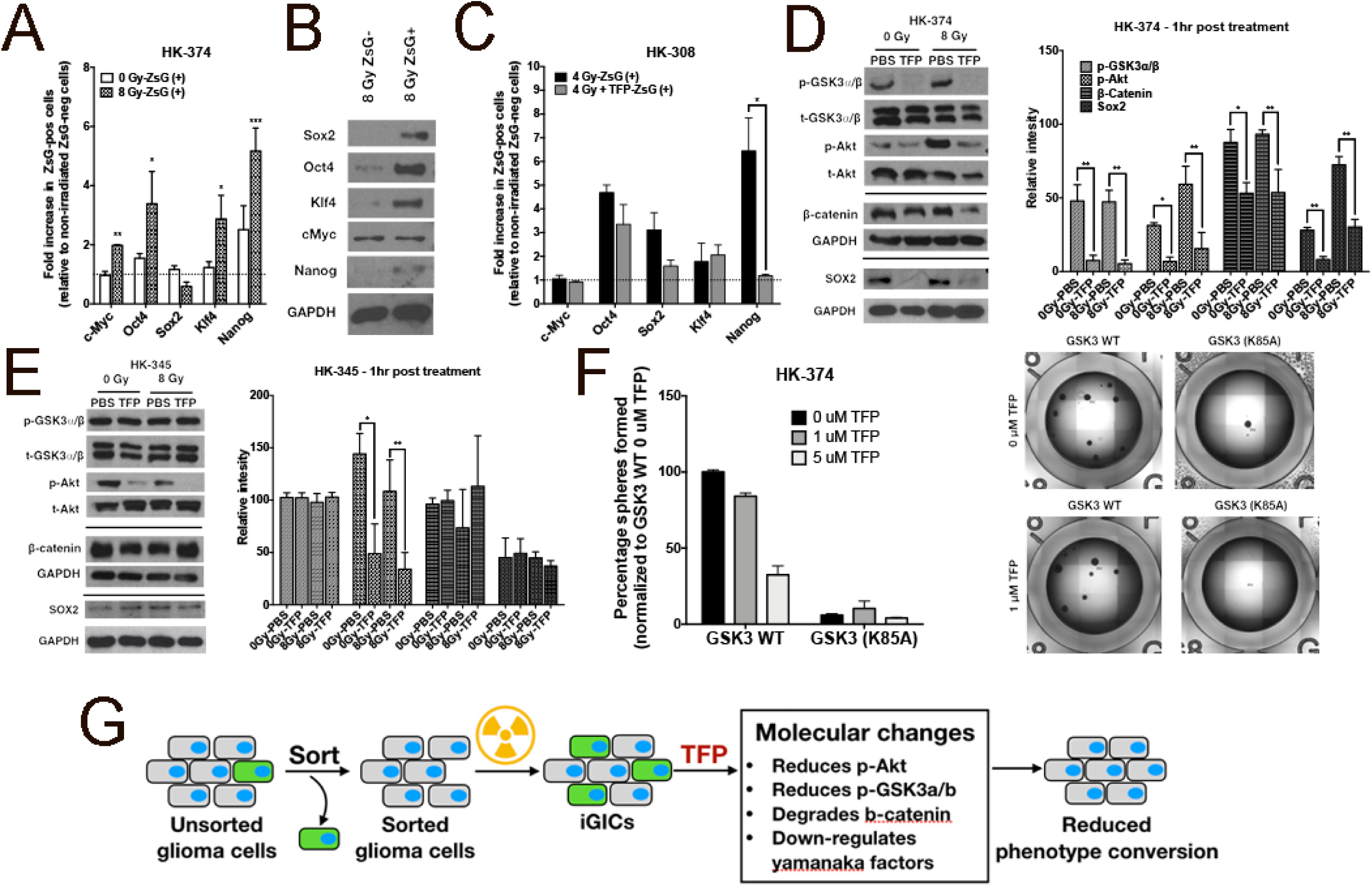
TFP reduces stem cell factors at the transcriptional and posttranslational level. ZsGreen-cODC-negative cells were collected after sorting the HK-374 ZsGreen-cODC vector expressing cells and plated in a 6-well plate at a density of 50,000 cells per well. The cells were irradiated at 0 or 8 Gy the following day. Five days after incubation, the cells were detached and re-sorted to collect both ZsGreen-cODC-positive and -negative cells. **(A)** qPCR was performed using RNA isolated from these samples. GAPDH was used as the internal control. The fold change in expression levels of Yamanaka factors in 0 Gy and 8 Gy ZsGreen-cODC-positive cells was calculated by 2^-ddCt method by normalizing it to non-irradiated ZsGreen-cODC negative cells. **(B)** Western blotting was performed with the proteins extracted from 8 Gy ZsGreen-negative and –positive cells and the membrane was blotted for Sox2, Oct4, Klf4, c-Myc and Nanog. GAPDH was used as the internal control. **(C)** ZsGreen-cODC-negative cells were collected after sorting the HK-308 ZsGreen-cODC vector expressing cells and plated in a 6-well plate at a density of 50,000 cells per well. The cells were pre-treated with TFP (10 uM) one hour before irradiating at 0 or 4 Gy the following day. Five days after incubation, the cells were detached and re-sorted to collect both ZsGreen-cODC-positive and -negative cells. qPCR was performed using RNA isolated from these samples. GAPDH was used as the internal control. The fold change in expression levels of Yamanaka factors in 0 Gy and 4 Gy ZsGreen-cODC-positive cells was calculated by 2^-ddCt method by normalizing it to non-irradiated ZsGreen-cODC negative cells. **(D & E)** 90% confluent plates with parent HK-374 and HK-345 cells were serum-starved overnight and pre-treated with either TFP (10 uM) or vehicle (Saline) for one hour. Just before irradiating these plates with a single dose of 0 or 8 Gy, second treatment with TFP or Saline was performed. Starting from the second treatment, the samples were collected at exactly one hour time point for protein analysis using Western blotting. The blots were analyzed for p-GSK3a/b, t-GSK3a/b, p-Akt, t-Akt, β-catenin, Sox2 and GAPDH. GAPDH was used as the loading control for β-catenin and Sox2. **(D & E)** The intensity of each band was calibrated using ImageJ and presented as density ratio of gene over GAPDH in HK-374 and HK-345 cells. **(F)** HK-374 cells were transduced with lenti-viral vector expressing GSK3 (K85A) mutant gene. HK-374 containing GSK3 WT cells along with HK-374 cells with GSK3 (K85) mutant gene were plated in 96-well plates and treated with TFP (0, 1 and 5 uM). This set up was maintained with regular feeding of the spheres with 10X growth factors. The number of spheres formed in each condition was counted and normalized against the GSK3 WT non-irradiated control. Representative images of GSK3 WT and GSK3 (K85A) mutant cells at 0 and 5 uM TFP. (**G**) Schematic representation on how TFP inhibits phenotype conversion. All experiments in this figure have been performed with at least 3 biological independent repeats. *p*-values were calculated using unpaired t-test. * *p*-value < 0.05, ** *p*-value < 0.01 and *** *p*-value < 0.001.

### TFP activates GSK3α and decreases β-catenin and Sox2 levels

GSK3 is a kinase that phosphorylates already phosphorylated proteins to mark them for proteasomal degradation thus, affecting cell signaling in multiple ways (31). Among others, downstream targets include β-catenin, PTEN, p65, HIF1, c-Myc and Oct4 (31). Active GSK3 lacks phosphorylation on Ser9 or Ser21 and counteracts a stem-cell phenotype by phosphorylating key proteins in developmental pathways, such as Oct4 and β-catenin (32), thereby targeting them for degradation. Therefore, inactive GSK3 (phosphorylated on Ser9/21) supports stemness by allowing stem cell factors to escape degradation. In addition, Akt signaling promotes stemness by leading to phosphorylation of Sox2 (33) and phosphorylation of Sox2 requires phosphorylated Akt (33). Phosphorylated Sox2 participates in a transcriptional protein complex composed of XPC, Oct4, Rad23B and Cent2 that is responsible for initiating transcription of the Nanog gene (34). Signaling through D_2_-like dopamine receptors leads to phosphorylation of Akt (35) and subsequent inhibitory GSK3 phosphorylation by Akt (36), thus supporting posttranslational stabilization of stem-cell factors. In order to understand how TFP interferes with the acquisition of a stem cell state, HK-374 cells were plated and serum-starved overnight and irradiated with 0 or 8 Gy in the presence or absence of TFP. TFP treatment led to a rapid loss of constitutive and radiation induced phosphorylation of Akt in HK-374 and HK-345 cells at the 1-hour time point. Loss of the inhibitory serine-21 phosphorylation of GSK3α was detected in HK-374 cells in which TFP prevented phenotype conversion. (**Figure 3D**). In contrast, phosphorylation of GSK3α remained unchanged in HK-345 cells (**Figure 3E**) in which TFP fails to prevent radiation-induced phenotype conversion (**Figure 2B**). Furthermore, TFP treatment reduced constitutive and radiation-induced β-catenin and Sox2 levels at 1 hour after TFP treatment in HK-374, but not in HK-345 cells (**Figure 3 D/E**).

To show dependence on GSK3, we next overexpressed a dominant-negative kinase-dead form (K85A) of GSK3-beta. Loss of GSK3-beta kinase activity significantly reduced the self-renewal capacity of HK-374 cells measured in a sphere-formation assay. This was expected given the multiple known targets of GSK3. Importantly, overexpression of the kinase-dead form of GSK3 rendered the cells unresponsive to TFP in this assay, thus further supporting the role of GSK3 in the response to TFP (**Figure 3F**). A graphical summary of these findings is presented in **Figure 3G**.

### Radiation induces developmental gene expression signatures in glioma cells

In order to uncover if radiation –aside from its DNA damaging effects and associated cellular responses– also affects developmental programs and to identify additional targets of TFP we next performed RNA-Seq to map gene expression changes induced in GBM cells by radiation. HK-374 ZsGreen-cODC-negative cells were sorted by FACS, treated with TFP and irradiated with 4 Gy. One and 48 hours after irradiation total RNA was extracted and subjected to RNASeq to study pathways immediately engaged by TFP treatment (1 hour) and their downstream consequences for the radiation response (48 hours).

Differential gene expression analysis revealed that 1-hour treatment with TFP induced changes in the expression levels of 884 genes (**Figure 4A**, left panel). In comparison, a single dose of 4 Gy led to 37 differentially expressed genes with the majority (29/37) of the genes down-regulated (**Figure 4A**, middle panel). Combination of TFP treatment with radiation changed the expression of 119 genes (**Figure 4A**, right panel). However, at one hour after drug treatment no genes were differentially expressed between cells treated with TFP and radiation versus radiation alone (**Figure 4A**, Venn Diagram). In contrary, 48 hours after drug treatment we identified 2952 differentially expressed genes when comparing cells treated with TFP and radiation versus radiation alone, creating a distinct gene expression profile in hierarchical clustering (**Figure 4B**). We selected the top 9 up-regulated and top 10 down-regulated genes in cells treated with TFP and radiation versus radiation alone and confirmed the expression changes using real-time qRT-PCR (**Figure 4C**).

**Figure 4.**
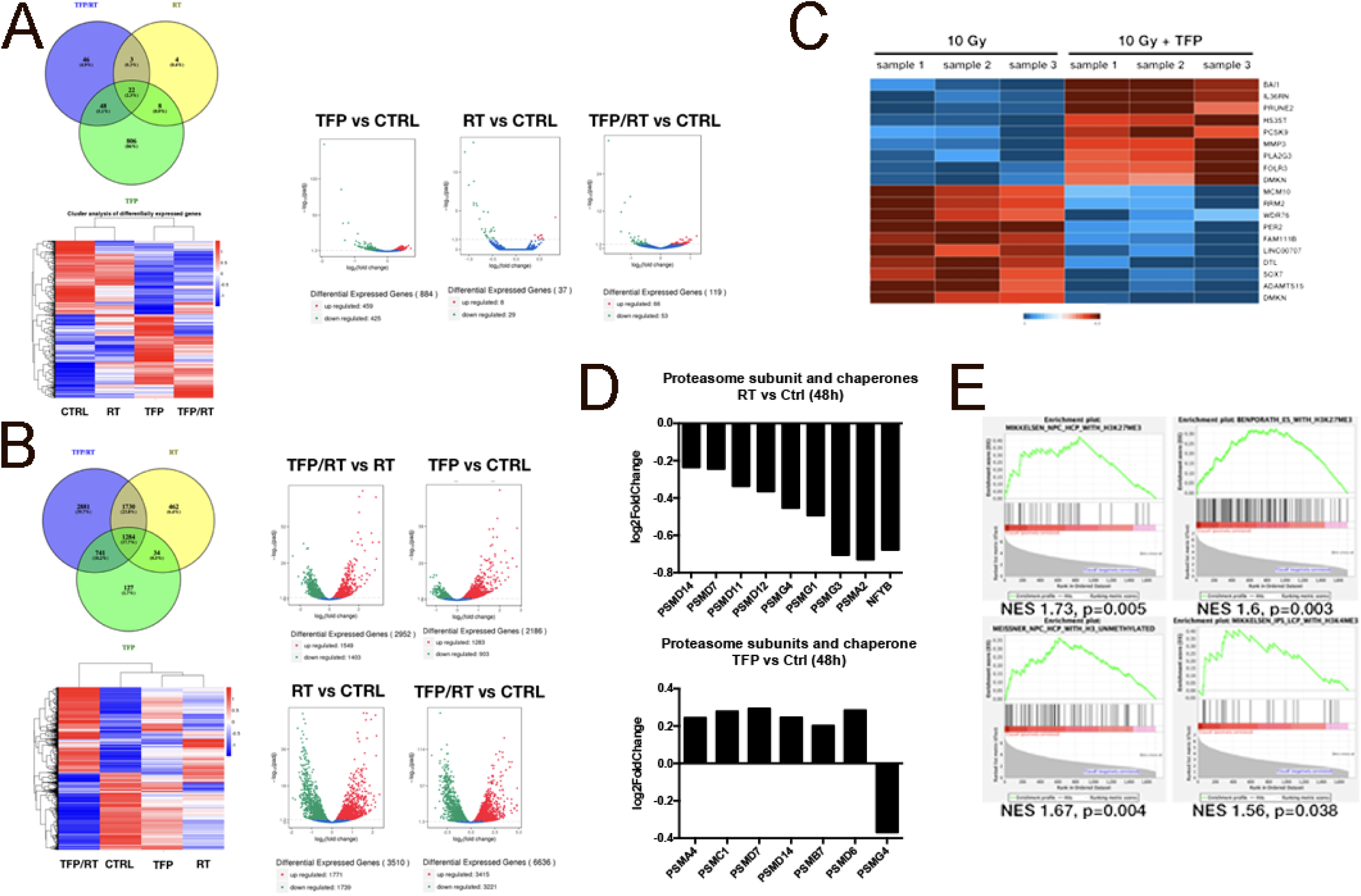
TFP interferes with DNA replication and cell cycle progression in GBM. **(A)** Venn diagram, volcano diagrams and heat map of sorted non-GICs from HK-374 cells, one hour after treatment with radiation (8Gy), TFP, or TFP and radiation compared to untreated control cells. (**B**) Venn diagram, volcano diagrams and heat map of sorted non-GICs from HK-374 cells, 48 hours after treatment with radiation, TFP, or TFP and radiation compared to untreated control cells. (**C)** Heat map showing the results of real-time RT-PCR for the top 9 up-regulated and top 10 down-regulated genes in HK-374 non-GICs comparing combination treatment of TFP and radiation with radiation alone, 48 hours after treatment. (**D**) Proteasome subunit and proteasome assembly chaperone mRNA expression 48 hours after radiation or TFP treatment. (**E**) Gene set enrichment analysis was performed on the data obtained by RNAseq analysis to identify differentially expressed genes that overlapped with curated gene sets representing expression of gene signatures pertaining to genetic and chemical perturbations.

In agreement with our previous studies (28), radiation led to the down-regulation of proteasome subunit and NF-YB expression at 48 hours after irradiation (**Figure 4D**, left panel). NF-YB is a subunit of the nuclear transcription factor Y (NF-Y) responsible for the concerted regulation of proteasome subunit expression (37). In contrary, TFP treatment increased proteasome subunit expression (**Figure 4D**, right panel). Expression of two inducible beta-type proteasome subunits, PSMB8 (1.68-fold) and PSMB9 (1.47-fold), was up-regulated in cells treated with radiation and TFP compared to radiation alone. These data supported our previous observations (11, 28) that loss of proteasome function is a major event in the acquisition of cancer stem cell traits after irradiation as it allows for posttranslational stabilization of stem cell factors, otherwise degraded by the proteasome.

In order to study if radiation engages developmental gene expression programs we performed a gene set enrichment analysis (38, 39) of our RNA-Seq data computing the overlap of differentially expressed genes with 3302 curated gene sets representing expression signatures of genetic and chemical perturbations.

At 48 hours after irradiation differentially expressed genes overlapped with gene sets found upregulated in neural stem progenitor cells, embryonic stem cells and induced pluripotent stem cells (**Figure 4E**).

### Epigenetic Changes in Glioma cells in response to radiation

Since we observed re-expression of Yamanaka transcription factors in response to radiation we next performed ChIP-PCR against the promoter regions of Sox2, Oct4 and Nanog and found radiation-induced acquisition of an open chromatin state for these promoters that peaked at 48 hours after irradiation, thus indicating radiation-induced epigenetic remodeling (**Figure 5A**). This was further supported by data obtained in a histone ELISA assay that revealed significant global changes in histone acetylation and methylation, 48 hours after irradiation with 4 Gy (**Figure 5B**). ATAC-Seq at the same 48 hour time point revealed acquisition of an open chromatin state in 1785 gene regions and a closed chromatin state in 3637 in irradiated cells versus unirradiated cells covering both, promoter and non-promoter regions of the genome (**Figure 5C/D**). A motif search analyzing the differentially opened and closed promoter regions revealed radiation-induced open chromatin with binding motifs for the developmental transcription factors Klf4, Pou3f2, E2F2, RUNX1 and MYC. Conversely, our motif search revealed a radiation-induced closed chromatin state in promoter regions with the binding motifs for Nkx3-1, a prostatic tumor suppressor, Pbx1 and Mtf1 involved in cell fate decisions, the androgen receptor binding motif and the binding motif for Smad4, a factor correlated with decreased survival in GBM when downregulated (**Figure 5E**).

**Figure 5.**
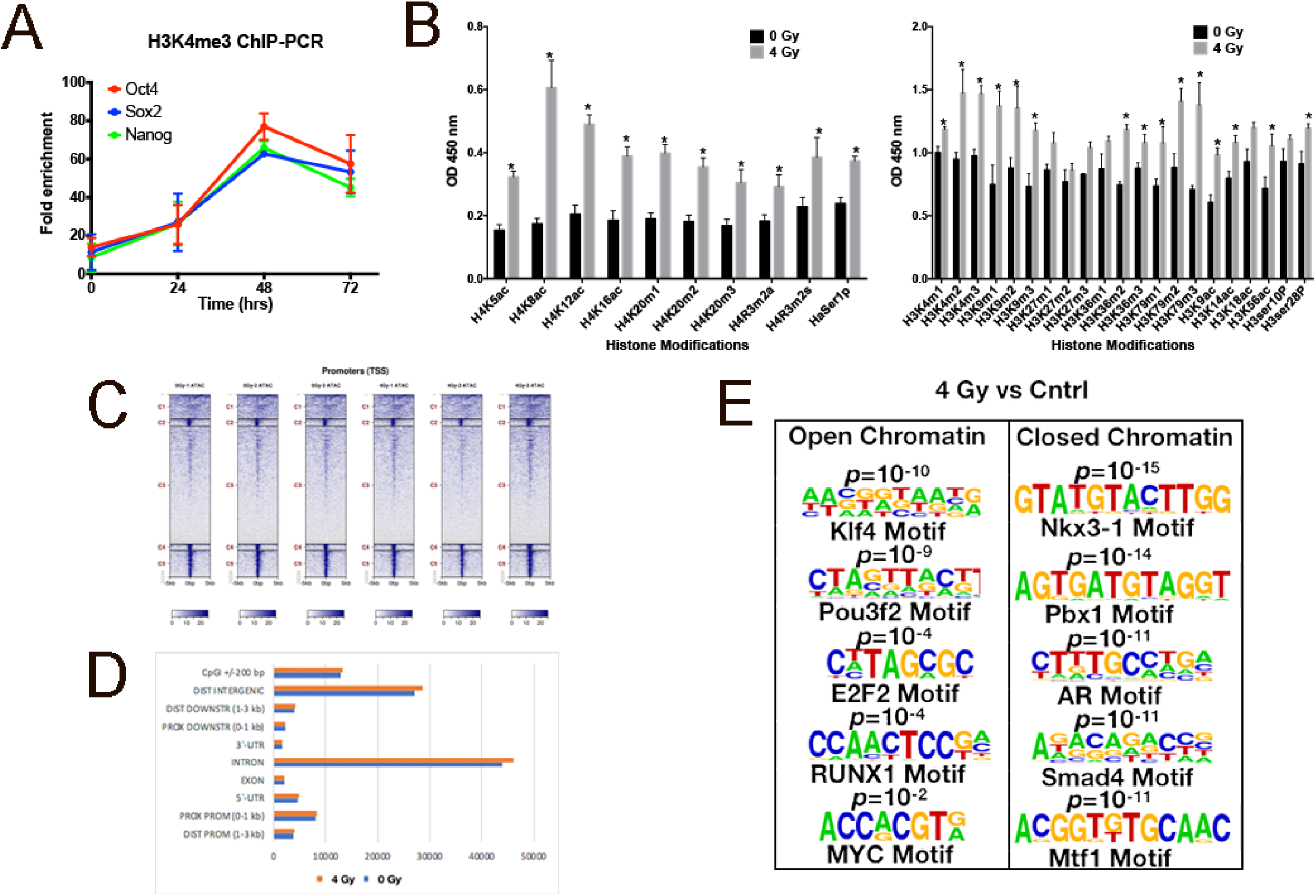
Radiation induces epigenetic changes in GBM. 70% confluent plates of HK-374 cell cultures were irradiated with 0 or 4 Gy and 48 hours after irradiation the cells were extracted for proteins and used to perform ChIP-PCR assay to analyze for H3K4me3 active chromatin interactions with specific proteins such as Sox2, Oct4 and Nanog **(A)**. The same set of proteins was also used to perform ELISA for H4 and H3 histone modifiers **(B)**. ATAC-seq samples were collected after 48 hours of 0 or 4 Gy irradiation in HK-374 cells. These samples were then analyzed for acquisition of an open chromatin and a closed chromatin state in 0 versus 4 Gy samples **(C and D)**. A motif search on both samples was also performed to identify the differentially opened or closed promoter regions with binding motifs for the developmental transcription factors such as Klf4, Pou3f2, E2F2, RUNX1 and MYC **(E)**.

### TFP delays tumor growth in murine models of GBM

Based on the *in vitro* effect of TFP on the sphere forming capacity of patient-derived glioma specimen (**Figure 2F**), we tested whether TFP also had anti-tumor activity in a murine model of GBM *in vivo*. GL261-Luc or GL261-StrawberryRed murine GBM cells, engineered to express firefly luciferase or StrawberryRed were implanted into the brains of C57Bl/6 mice. On day 7 post tumor implantation, baseline bioluminescence pictures were acquired and animals were randomized and treated with either saline, 20 mg/kg TFP, radiation (10 Gy), or a combination. Mice bearing StrawberryRed-expressing GL261 tumors were treated with only saline or TFP starting on day 7. The TFP dose of 20 mg/kg, which has previously been reported as the MTD in mice (23) was well tolerated in our experiments. After an initial period of TFP-induced sedation that lasted 2-3 hours, the mice did not show any side effects and TFP-treated animals did not show the tumor associated weight loss that was observed in the control group (**Supplementary Figure 2**).

Mice bearing StrawberryRed-expressing tumors were sacrificed at different time points after TFP treatments. The brains were explanted, digested and the number of StrawberryRed-positive cells was quantified and assayed for self-renewal capacity, a functional assay for GICs. *In vivo* TFP treatment led to a decrease in the total number of tumor cells per brain in the mouse GL261 GBM model and a significant decrease in the total number of tumor cells in human HK-374 and HK-157 PDOXs (**Figure 6A).** Importantly, TFP treatment reduced the sphere formation capacity of the surviving tumor cells (**Figure 6B**).

**Figure 6.**
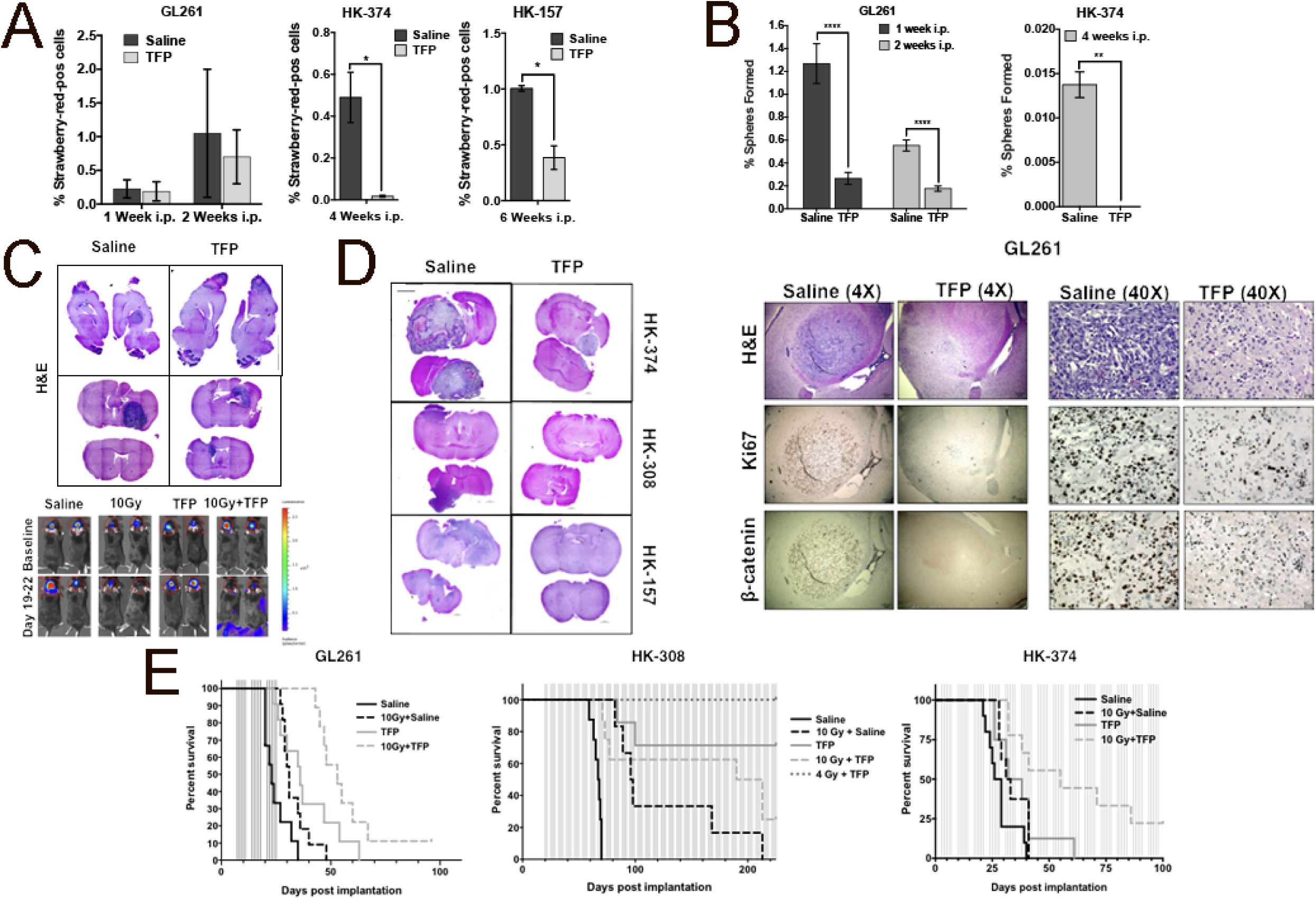
TFP in combination with radiation prolongs survival in mouse models of GBM. **(A)** 2×10^5^ GL261-StrawberryRed cells were implanted into the brains of the C57BL/6 mice and 3×10^5^ HK-374 and HK-157-StrawberryRed cells were implanted into the brains of NSG mice. Tumors were grown for 3 days (HK-374) and 7 days (HK-157 and GL261) for successful grafting. Mice bearing tumors were injected intra-peritoneally on a 5-day on / 2-day off schedule for 1 or 2 weeks (GL261), 4 weeks (HK-374) and 6 weeks (HK-157) either with TFP (20 mg/kg) or saline. At the end of the treatment, brains of these mice were extracted, tumors digested and sorted for StrawberryRed cells. The data from the sort were plotted and presented as percentage of StrawberryRed-positive cells of the total number of cells in each GBM line. **(B)** The sorted StrawberryRed cells from (A) were plated on 96-well plates in serum-free medium and fed with fresh growth factors every 2-3 days. The number of spheres formed in each condition was counted and normalized against the respective control. **(C)** H&E stained sagittal and coronal sections of the brains of C57BL/6 mice intra-cranially implanted with 2×10^5^ GL261 cells. Tumors were grown for 7 days for successful grafting. Mice were injected with TFP or saline intra-peritoneally on a 5-day on / 2-day off schedule for 3 weeks. TFP treatment led to a marked reduction in tumor size. **(D)** H&E stained coronal sections of the NSG mice brains implanted with HK-374, HK-308 and HK-157-GFP-Luc cells which were treated continuously with either TFP (sub-cutaneous, 20 mg/kg) or saline until they met the criteria for study endpoint. **(E)** 4X and 40X images of coronal sections of brains from C57BL/6 mice implanted with GL261-GFP-Luc cells intracranially and treated with either TFP (20 mg/kg) or saline intraperitoneally for 2 weeks and stained for H&E, Ki67 and β-catenin. **(F)** Survival curves for C57BL/6 mice implanted intra-cranially with 2×10^5^ GL261-GFP-Luc cells and NSG mice injected intra-cranially with 3×10^5^ HK-308-Luc or HK-374-GFP-Luc cells. Tumors were grown for 3 days (HK-374), 7 days (GL261) and 21 days (HK-308) for successful grafting. Mice were injected with either TFP (20 mg/kg) or saline intra-peritoneally for GL261 for three weeks, and sub-cutaneously for HK-308 and HK-374 lines, continuously until they reached the study endpoint. Log-rank (Mantel-Cox) test for comparison of Kaplan-Meier survival curves indicated significant differences in the TFP treated mice compared to their respective controls. *p*-values in GL261 study: saline vs. TFP (*** *p*-value = 0.00014), saline vs. 10 Gy + saline (* *p*-value = 0.0126), saline vs. 10 Gy + TFP (*** *p*-value <0.0002), TFP vs. 10 Gy (*p*-value = 0.5320), TFP vs. 10 Gy + TFP (* *p*-value = 0.0324). *p*-values in HK-308 study: saline vs. TFP (*** *p*-value = 0.00014), saline vs. 10 Gy + saline (*** *p*-value = 0.0004), saline vs. 10 Gy + TFP (**** *p*-value ≤ 0.0001), saline vs. 4 Gy + TFP (** *p*-value = 0.0011), TFP vs. 10 Gy (* *p*-value = 0.0141), TFP vs. 10 Gy + TFP (*p*-value = 0.1052), TFP vs. 4 Gy + TFP (*p*-value = 0.2137). *p*-values in HK-374 study: saline vs. TFP (\**p*-value = 0.0361), saline vs. 10 Gy + saline (* *p*-value = 0.0413), saline vs. 10 Gy + TFP (*** *p*-value = 0.00025), TFP vs. 10 Gy (*p*-value = 0.7217), TFP vs. 10 Gy + TFP (\**p*-value = 0.0273). GBM-StrawberryRed experiments in this figure have been performed with at least 2 biological repeats. *p*-values were calculated using unpaired t-test. * *p*-value < 0.05, ** *p*-value < 0.01 and **** *p*-value < 0.0001.

The loss in total tumor cell numbers and reduction in sphere-forming capacity was accompanied by a reduction in tumor size in tissue sections in GL261 (**Figure 6C**), HK-157, HK-308 and HK-374 (**Figure 6D**) tumor-bearing mice.

To confirm the effects of TFP on the Wnt pathway *in vivo*, GL261 cells were implanted into the brains of C57BL/6 mice. After grafting, the animals were administered 5 daily injections of TFP for two weeks. Tissue sections of the brains revealed a reduction in tumor size and loss of Ki67^+^ and β-catenin^+^ cancer cells (**Figure 6E**), which was in line with our *in vitro* observations (**Figure 3**).

Consistent with these observations, the median survival in GL261-bearing animals treated with TFP was 36 days compared to 23 days in the control group of animals (*p* = 0.00014). Irradiation with a single dose of 10 Gy increased the median survival from 23 to 31 days (*p* = 0.0126). Compared to saline-treated animals, a combination of a single dose of 10 Gy with TFP treatment significantly extended the median survival to 53 days (*p* = 0.0002) (**Figure 6F**, left panel).

In order to assess the effect of TFP on human GBM models we implanted two different human GBM specimens, the HK-308, a slow growing specimen and the HK-374, a fast-growing specimen, into the brains of NSG mice. Tumor take in HK-308-implanted mice was confirmed by bioluminescence imaging and the animals were randomized into the different treatment groups. Mice were treated with 5 weekly TFP (s.c.) injections or saline until they reached endpoints for euthanasia. Radiation treatment with a single dose of 10 Gy increased median survival to 97 days compared to 67.7 days in the control group (*p* = 0.0004). Remarkably, the combining 10 Gy with TFP extended survival to > 200 days (*p* < 0.0001). Combining TFP with a reduced 4 Gy dose of radiation resulted in 100 % of the animals surviving > 200 days post implantation (**Figure 6F**, middle panel). The observation that the combined treatment with TFP and 10 Gy was inferior to TFP treatment alone could reflect CNS toxicity after 10 Gy in NSG mice, which lack non-homologous end-joining DNA repair.

Median survival in animals implanted with the more aggressive HK-374 specimen was 27.5 days and treatment with TFP or 10 Gy extended the median survival to 32 (*p* = 0.0361) or 35 days (*p* = 0.0413) respectively. Animals treated with the combination of TFP and 10 Gy had a median survival of 55 days (*p* = 0.00025) (**Figure 6F**, right panel).

## Discussion

Despite decades of research aimed at improving treatment outcome for patients suffering from GBM the survival rates for this disease remain dismal. The reasons for the failure of RT to control GBM are multiple and include the highly infiltrative nature of GBM (40), which makes complete resection of the tumor nearly impossible, large areas of tumor hypoxia (41) that diminish the efficacy of ionizing radiation (42) and many chemotherapeutic agents (43), and the intrinsic resistance of glioma-initiating cells against established anti-cancer therapies (9, 10). Yet, although radiation therapy does not provide cure in the GBM setting it remains one of the few therapies that significantly prolongs the survival of GBM patients (3, 44).

The underlying biology of fractionated radiation therapy has been elegantly summarized by Rodney Withers in the 4 Rs of RT. In this model, redistribution within the cell cycle and reoxygenation of hypoxic tumor regions between fractions lead to therapeutic gain, whereas repopulation of the tumor by surviving cells and repair of DNA damage can lead to therapeutic loss (45). The intrinsic radiation sensitivity has been proposed as a 5^th^ R of RT (46) however, this concept is currently less widely accepted, mainly because it will be co-determined by the tumor oxygenation status, position of cancer cells in the cell cycle and their ability to repair DNA damage, which altogether might exaggerate or diminish the intrinsic radiosensitivity that is determined genetically and/or epigenetically.

The 4 (5) Rs of RT consider intra-tumoral heterogeneity to some extent but if interpreted in the context of the cancer stem cell hypothesis and thus, intra-tumoral hierarchy, the model assumes that the differentiation of cancer stem cells during repopulation is unidirectional. Even if one takes into consideration that cancer stem cells are exceptionally radioresistant but rare, this assumption commands that any given tumor can be cured by radiation dose escalation and the mainstream of research in clinical radiation oncology seeks to enable this through increased precision in dose delivery and reduction in normal tissue dose. However, the infiltrative nature of GBM together with the normal tissue radiation tolerance of the brain limit the success of radiation therapy against GBM to the *status quo*.

The phenotype conversion of non-tumorigenic cells into radio-resistant GICs through micro-environmental factors such as low pH and hypoxia (12, 47) or through radiation-induced epigenetic changes described in our present study potentially adds reprogramming as a 6^th^ R to the radiobiology of cancer. Our observations that this phenomenon occurs spontaneously but is even more pronounced after irradiation, the fact that it is not restricted to a population of cells prone to phenotype conversion, and the increased self-renewal capacity and radiation resistance of induced GICs suggest that it potentially has significant clinical relevance in that the ratio of phenotype conversion to cell killing of existing tumor-initiating cells per fraction determines local control of the tumor. This distinction is highly relevant because it suggests that unless a therapy given concurrently with radiation interferes with this phenomenon, radiation dose escalation will unlikely lead to local tumor control as long as the rate of phenotype conversion outpaces cell killing of existing tumor-initiating cells.

While generally thought to be associated with neuronal signaling, recent studies have supported a role for the dopamine D2 receptor in gliomagenesis (48, 49). Importantly, in the latter study dopamine D2 receptor expression was increased in response to temozolomide treatment, elevated in CD133-positive cells, promoted self-renewal and led to activation of hypoxia-inducible factors 1 and 2 under normoxic conditions (49).

Here we present the dopamine receptor antagonist, TFP as an already FDA-approved, blood-brain barrier-penetrating drug that not only targets preexisting glioma-initiating cells but also interferes with the process of phenotype conversion of GBM cells into radio-resistant induced glioma-initiating cells.

Short-term 21-day monotherapy with lower doses (10 mg/kg) of TFP against U87MG xenografts has been previously reported but did not yield a survival benefit (50). A second study reported radiosensitization of U87MG and U251 cells based on increased gamma-H2AX foci formation and inhibition of DNA repair by homologous recombination. In a single survival experiment using a patient-derived specimen, animals were treated with 3 × 5 Gy and TFP at 1 mg/kg. No information about radiation dosimetry was given and although the total radiation dose of 15 Gy was substantial, it showed little effect by itself (51).

Our study suggests that ionizing radiation elevates differentiated GBM cells into a state that allows for acquisition of cancer stem cell traits and that this process can also occur spontaneously although at a much lower frequency. The observation that radiation leads to epigenetic remodeling characterized by an open chromatin state in the promoter region of genes involved in the reprogramming of somatic cells into induced pluripotent stem cells and the promoter regions of their target genes consequently resulting in the activation of developmental gene expression programs indicates the existence of potential new targets to increase the efficacy of radiation therapy.

Mechanistically, we were able to connect pathways known to be affected by TFP treatment like Akt signaling to the posttranslational regulation of key molecular mediators of stem cell traits in cancer, like Sox2 and β-catenin. However, our RNASeq analysis showed that responses to TFP in combination with radiation are complex and target DNA biosynthesis, cell cycle progression and cell division in GBM cells.

Although we found that TFP is not a classical radio-sensitizer in that it does not affect the intrinsic radiosensitivity of GICs directly, it significantly prolongs the survival in syngeneic and PDOX mouse models of glioma when combined with RT. It is noteworthy that NSG mice are considered radiosensitive with an LD50 in total body irradiation experiments of 6.5 Gy (52). However, it is known that when tumors are irradiated in mice carrying the *Scid* mutation, the radiosensitivity of the microenvironment does not alter tumor responses (53). Furthermore, 10 Gy radiation doses to the mostly post-mitotic midbrain were well tolerated in our study and animals survived for over 200 days without signs of radiation toxicity, thus supporting that NSG mice can be used for radiation studies of GBM PDOXs.

We conclude that a combined treatment with the dopamine receptor antagonist TFP and radiation could be an effective new treatment against GBM as it not only targets GBM cells but prevents the acquisition of a more aggressive phenotype of GBM cells induced by radiation. The data reported in our study indicate the need for continuous drug treatment and the necessity of dose-finding studies for future combination trials, ideally with second generation dopamine receptor antagonists that exhibit a more favorable side effect profile than TFP.

## Acknowledgements

FP was supported by grants from the *National Cancer Institute* (CA137110, CA161294). MP, PLN, TC, LL, HK, and FP were supported by a grant from the *National Cancer Institute* (UCLA Brain Tumor SPORE P50CA211015).

