## Supplementary Material for "The Dopamine Receptor Antagonist TFP Prevents Phenotype Conversion and Improves Survival in Mouse Models of Glioblastoma"

**This PDF file includes:**

Supplementary text

Supplementary Table1

Supplementary figures 1 & 2

**Material and Methods**

*Senescence analysis*

HK-374 ZsGreen-cODC expressing cells were plated in 6 well plates at a density of 50,000 cells/well and the next day irradiated with 0 or 8 Gy. 5 days after irradiation the plates were assayed for senescence via the Senescence β-galactosidase Cell Staining Kit (Cell Signaling, Catalog # 9860) and the staining was performed as per manufacturer’s protocol. Briefly, the plates were rinsed with PBS and 1 ml of the 1X Fixative Solution was added and incubated at room temperature for 10 minutes. The plates were rinsed twice with 1X PBS and stained with 1 ml of β-galactosidase staining solution (930 μl of 1X staining solution, 10 μl of 100X solution A, 10 μl of 100X solution B, 50 μl of 20 mg/ml X-gal stock solution) in each well. The plate was incubated at 37°C for 18 h in a dry incubator (no CO_2_). The color development in the senescent cells and fluorescence in the ZsGreen-positive cells were captured at 20X using a digital fluorescence microscope (BZ-9000, Keyence, Itasca, IL).

*Brain tissue digestion and flow cytometry*

2x10^5^ GL261-StrawberryRed and 3x10^5^ HK-374- or HK-157-StrawberryRed cells were implanted into the brains of the C57BL/6 or NSG mice respectively, as described above. Tumors were grown for 3 (HK-374) and 7 days (HK-157 and GL261) for successful grafting. Mice-bearing tumors were injected intra-peritoneally on a 5-days on / 2-days off schedule for 1 or 2 weeks (GL261), 4 weeks (HK-374) and 6 weeks (HK-157) either with TFP or saline. TFP was dissolved in sterile saline at a concentration of 2.5 mg/mL. All animals were treated with 20 mg/kg TFP. At the indicated time points after implantation the mice were sacrificed and tumor-bearing brains were dissected for further analysis.

For brain tumor dissociation, Miltenyi C tubes (gentleMACS C tubes, Cat # 130-093-237, Auburn, CA, USA) were initially prepped with 2X enzymes – 100 μL enzyme D, 50 μL enzyme R and 12.5 μL enzyme A provided in the mouse Tumor Dissociation Kit (Cat # 130-096-730, Miltenyi, Auburn, CA, USA) in 2.35 mL of DMEM/F12 media. The quadrant of the brain surrounding the site of tumor implantation was dissected and finely chopped with a scalpel. The minced tissue was then transferred to the C tubes and tumor pieces were further dissociated by running the m_impTumor_02 program once on the gentleMACS Dissociator (Cat # 130-093-235, Miltenyi, Auburn, CA, USA). The tubes were next placed in a shaker incubator at 37°C for 40 mins. After incubation, the tubes were once again placed in the gentleMACS Dissociator and subjected twice to the same dissociation program as above. The tubes were briefly centrifuged at 300 x g to collect the digested cells at the bottom of the tube. The cells were re-suspended in the enzyme-containing medium and filtered through a 70 μM filter into a 50 mL conical tube. The filter was washed with 10 mL DMEM/F12 media and centrifuged at 300 x g for 7 mins. The supernatant was aspirated and the pellet was re-suspended in 2 mL of ACK lysis buffer (Cat # 10-548E, BioWhittaker, Walkersville, MD, USA) to lyse the red blood cells. After 2 minutes, the conical tubes were centrifuged at 300 x g for 7 minutes and the supernatant was aspirated. The cell pellet was then re-suspended in 1-2 mL of GBM serum-free media. The cell suspension was sorted on the ARIAIII High-Speed Cell Sorter (BD Biosciences, San Jose, CA, USA) for StrawberryRed-expressing GBM cells directly onto a 96-well plate to assess self-renewal capacity of the cells in a functional assay. For flow cytometric analysis, the cells obtained from dissociating tumors from parent GBM cells were used as a negative control. Samples from TFP or saline treated tumors were gated against the negative control and the StrawberryRed-positive cells were graphed as percentage of StrawberryRed-pos cells.

For radiation-induced reprogramming experiments *in vivo* with GL261-StrawberryRed and GL261-BFP cells, GL261-StrawberryRed were first stained with an anti-Prominin-APC antibody (Miltenyi, Cat # 130-102-197) and the Prominin-positive cells were removed by FACS. The Prominin-negative cells were seeded into 6-well plates at a density of 50,000 cells/well. Cells were irradiated with 0 or 4 Gy and incubated at 37°C in a CO_2_ incubator for five days. After 5 days, GL261-StrawberryRed cells along with GL261-BFP cells were stained with an anti-Prominin-APC antibody to determine the percentage of prominin-positive cells in the total population. 25,000 cells from each cell population (GL261-StrawberryRed and GL261-BFP) and from each group (0 Gy or 4 Gy) were mixed together and a total of 50,000 cells were injected into mice intracranially. The mice were maintained for 3 weeks and after that the brains were dissociated as described above for flow cytometric analysis with GL261-StrawberryRed and GL261-BFP cells stained with anti-Prominin antibody.

*Quantitative Reverse Transcription-PCR*

Total RNA was isolated using TRIZOL Reagent (Invitrogen). cDNA synthesis was carried out using the SuperScript Reverse Transcription III (Invitrogen). Quantitative PCR was performed in the My iQ thermal cycler (Bio-Rad, Hercules, CA) using the 2x iQ SYBR Green Supermix (Bio-Rad). *C*_t_ for each gene was determined after normalization to GAPDH or RPLP0 and ΔΔ*C*_t_ was calculated relative to the designated reference sample. Gene expression values were then set equal to 2^−ΔΔCt^ as described by the manufacturer of the kit (Applied Biosystems). All PCR primers were synthesized by Invitrogen and designed for the human sequences of Yamanaka factors (Oct4, Sox2, Nanog, Klf4, c-Myc), dopamine receptors (DRD1, 2, 3, 4, 5), genes for RNASeq result validation (PDK4, ALDOC, SLC2A5, PDK2, ALDH2, PC, GFPT1, PFKFB4, GFPT2, TOMM22, MRPS27, SLC25A24, ME2, TIMM13, MRPL3, MRPL22, MRPS34, MRPL18, SLC25A5, MPC1, GPD2, TIMM21, CMC2, TIMM9) and PPIA, TBP and IPO8 as housekeeping genes (for primer sequences see supplement).

*Chromatin Immune Precipitation Polymerase Chain Reaction (ChIP-PCR)*

ChIP-PCR was performed using SimpleChIP^®^ Plus Enzymatic Chromatin IP Kit (Cell Signaling, Cat # 9005) by following manufacturer’s protocol. Briefly, HK-374 cells were plated in 15 cm dishes and cultured until they reached confluence. The plates were irradiated at 0 or 4 Gy and 48 hours later the cells were analyzed. To crosslink proteins to DNA, 540 μl of 37% formaldehyde was added to each dish containing 20 ml of culture medium. The dish was swirled briefly and incubated at room temperature for 10 min. 2 ml of 10X glycine was added to each dish, swirled briefly and then incubated at room temperature for 5 minutes. The cells were next washed with ice-cold PBS. 2 ml of ice-cold PBS and Protease Inhibitor Cocktail (PIC) was added to the dish and the cells were scraped into the cold buffer. Cells were centrifuged at 2,000 x g for 5 min at 4°C. The supernatant was discarded and the cells were re-suspended in 1 ml of ice-cold 1X Buffer A + PIC + DTT per IP prep and incubated on ice for 10 minutes. The cell nuclei were pelleted by centrifuging at 2,000 x g for 5 min at 4°C. The supernatant was discarded and the pellet was resuspended in 100 μl of Buffer B + PIC + DTT per IP prep. 0.5 μl of micrococcal nuclease (Cell Signaling, Cat # 10011) per IP prep was added, the tubes inverted several times and incubated for 20 mins at 37°C. Digestion was stopped by adding 10 μl of 0.5 M EDTA (Cell Signaling, Cat # 7011) per IP and the tubes were placed on ice for 2 mins. The nuclei were centrifuged at 16,000 x g for 1 min at 4°C. The pellet was resuspended in 100 μl of 1X ChIP Buffer + PIC per IP and incubated on ice for 10 min. The samples were then sonicated with several pulses to break the nuclear membrane. The samples were incubated on ice for 30 sec between pulses. The lysates were centrifuged at 9,400 x g for 10 minutes at 4°C. Next, chromatin immunoprecipitation was performed using the cross-linked chromatin samples. For each immunoprecipitation, 2 μl of the immunoprecipitating antibody (Sox2 (Cat # 3579), Oct4 (Cat #2750), Nanog (Cat # 4903), Cell Signaling) were added to 500 μl of the diluted chromatin. The samples were rotated overnight at 4°C. The next day, 30 μl of the ChIP-Grade Protein G Magnetic Beads (Cell Signaling, Cat # 9006) was added to each IP sample and incubated at 4°C for 2 hours. The suspensions were placed on a magnetic separation rack (Cell Signaling, Cat # 7017) to pellet the magnetic beads and bound protein. The samples were washed, 150 μl of the 1X ChIP Elution Buffer was added and incubated at 65°C for 30 minutes. The magnetic beads were removed using the separation rack. The eluted chromatin was then transferred to a fresh tube. To remove the cross-links, 6 μl of 5 M NaCl and 2 μl of Proteinase K (Cell Signaling, Cat # 10012) were added and the samples incubated at 65°C for 2 hours. The eluted chromatin was used to perform quantitative PCR by using control primers included in the kit for the human RPL30 Exon 3 primers (Cell Signaling, Cat #7014). The primers used for this study were SimpleChIP^℗^ Human Sox2 Promoter primers (Cell Signaling, Cat # 4649S), SimpleChIP^℗^ Human Oct4 Promoter primers (Cell Signaling, Cat # 4641S), SimpleChIP^℗^ Human Nanog Promoter primers (Cell Signaling, Cat # 95064S).

*Histone Enzyme-Linked Immunosorbent Assay*

HK-374 cells were irradiated with 0 and 4 Gy. Fourty-eight hours after irradiation histone proteins were isolated using EpiQuik Total Histone Extraxtion Kit (Epigentek, Farmingdale, NY, Cat #OP-0006). Briefly, cells were harvested by trypsinizing and then centrifuging at 1000 rpm for 5 min at 4°C. The cells were re-suspended in 1X Pre-Lysis Buffer and incubated on ice for 10 min with gentle stirring. The suspension was then centrifuged at 3000 rpm for 5 min at 4°C. The pellet was re-suspended in 200 μL Lysis Buffer and incubated on ice for 30 min. The lysate was centrifuged at 12,000 rpm for 5 min at 4°C. The protein concentration was calibrated by BCA assay (Thermo Scientific™ Pierce™ BCA™, Waltham, MA, Cat #PI23227).

100 ng of the protein was used from each condition to analyze for Histone H3 and H4 modifications. ELISA for H3 (Epigentek, Cat # P-3100) and H4 (Epigentek, Cat # P-3102) histones was performed as per manufacturer’s protocol. Briefly, 49 μl of Antibody Buffer was added to all wells. For control wells 1 μl of Diluted Assay Control Protein was added to each standard well (5 and 25 ng/well). For sample wells, 4 μl of histone extracts was added. The plates were sealed and incubated at 37°C for 2 hours. The wells were rinsed thrice with 150 μl of the 1X Wash Buffer. 50 μl of the 1X Detection Antibody was added to each well, and incubated at room temperature for 1 hour. The wells were washed thrice and 100 μl of Developer solution was added to each well and incubated at room temperature for 5 min in the dark, after which 100 μl of Stop Solution was added to each well. The absorbance was read on a spectrophotometer (SpectraMax M5, Molecular Devices ) at 450 nm.

*RNASeq*

One and 48 hours after 4 Gy irradiation or sham irradiation, RNA was extracted from ZsGreen-cODC-negative HK-374 non-GICs using Trizol. RNASeq analysis was performed by Novogene (Chula Vista, CA). Quality and integrity of total RNA was controlled on Agilent Technologies 2100 Bioanalyzer (Agilent Technologies; Waldbronn, Germany). The RNA sequencing library was generated using NEBNext® Ultra RNA Library Prep Kit (New England Biolabs) according to manufacturer’s protocols. The library concentration was quantified using a Qubit 3.0 fluorometer (Life Technologies), and then diluted to 1 ng/uL before checking insert size on an Agilent Technologies 2100 Bioanalyzer (Agilent Technologies; Waldbronn, Germany) and quantifying to greater accuracy by quantitative Q-PCR (library molarity >2 nM). The library was sequenced on Illumina NovaSeq6000 with an average of 20M reads per RNA sample.

Downstream analysis was performed using a combination of programs including STAR, HTseq, and Cufflink. Alignments were parsed using the program Tophat and differential expressions were determined through DESeq2. Reference genome and gene model annotation files were downloaded from genome website browser (NCBI/UCSC/Ensembl) directly. Indexes of the reference genome were built using STAR and paired-end clean reads were aligned to the reference genome, using STAR (v2.5). HTSeq v0.6.1 was used to count the read numbers mapped of each gene. The FPKM of each gene was calculated based on the length of the gene and reads count mapped to this gene.

Differential expression analysis between irradiated and control samples (three biological replicates per condition) was performed using the DESeq2 R package (2_1.6.3). The resulting *p*-values were adjusted using the Benjamini and Hochberg’s approach for controlling the False Discovery Rate (FDR). Genes with an adjusted *p*-value of <0.05 found by DESeq2 were assigned as differentially expressed. To identify the correlations between differentially expressed genes, we generated heatmaps using the hierarchical clustering distance method with the function of heatmap, SOM (Self-organization mapping) and k-means using the silhouette coefficient to adapt the optimal classification with default parameters in R. Gene set enrichment analysis of differentially expressed genes was performed using the GSEA tool (version 4.0.1). Overlapping gene set with corrected *p*-values less than 0.05 were considered significantly enriched by differential expressed genes.

*siRNA Treatment*

70% confluent HK-374 cell cultures were grown in antibiotics-free culture media overnight. The next day the culture media was replaced with 1 ml of Opti-MEM™ reduced serum media. To this control or DRD2 siRNA-lipid complex (1:1) was added drop-wise and mixed well. Control and DRD2 siRNA was obtained from Dharmacon (Lafayette, CO). Lipofectamine® RNAiMAX transfecting reagent was obtained from Life technologies (Carlsbad, CA). The set up was then incubated at 37°C in a CO_2_ incubator for four hours. After transfection, the media was replaced with fresh culture media with 10% FBS. Seventy-two hours post siRNA transfection, the cells were serum starved for four hours and treated with 10 μM TFP for one hour. Proteins from these cells were lysed using RIPA buffer and protein concentrations were calibrated using BCA assay. Western blotting was performed to confirm DRD2 knockdown using antibodies against DRD2 (Millipore, Sigma, Cat # AB5084P, 1:1000) and to analyze its effect on p-GSK3α/β, t-GSK3α/β, β-catenin. GAPDH was used as the loading control. ImageJ software was used to perform the densitometry.

*Western Blotting*

GBM cells were serum starved overnight and the following day pre-treated with 10 μM TFP for one hour. Pre-treated HK-374 and HK-345 cells were irradiated with a single dose of 8 Gy immediately after a second treatment with 10 μM TFP. One hour after irradiation, the cells were lysed in 150 μl of ice-cold RIPA lysis buffer (10 mM Tris-HCl (pH 8.0), 1 mM EDTA, 1 % Triton X-100, 0.1 % Sodium Deoxycholate, 0.1 % SDS, 140 mM NaCl, 1 mM PMSF) containing proteinase inhibitor (Thermo Fisher Scientific) and phosphatase inhibitor (Thermo Fisher Scientific). The protein concentration in each sample was determined by BCA protein assay (Therma Fisher Scientific) and samples were denaturated in 4X Laemmli sample buffer (Bio-Rad) containing 10% β-mercaptoethanol for 10 minutes at 95°C. Equal amounts of protein were loaded onto 10% SDS-PAGE gels (1X Stacking buffer - 1.0 M Tris-HCl, 0.1% SDS, pH 6.8, 1X Separating buffer - 1.5 M Tris-HCl, 0.4% SDS, pH 8.8) and were subjected to electrophoresis in 1X Running buffer (12.5 mM Tris-base, 100 mM Glycine, 0.05% SDS), initially at 40 V for 30 minutes followed by 80 V for two hours. Samples were then transferred onto 0.45 μM nitrocellulose membrane (Bio-Rad) for two hours at 80 V. Membranes were blocked in 1X TBST (20 mM Tris-base, 150 mM NaCl, 0.2% Tween-20) containing 5% milk or 5% bovine serum albumin (BSA) for 20 minutes and then washed with 1X TBST followed by incubation with primary antibodies against p-Akt (Cat # 9271S, 1:1,000, Cell Signaling), p-GSK3α/β (Cat # 9327S, 1:1000, Cell Signaling), β-catenin (Cat # 8480S, 1:1,000, Cell Signaling), GAPDH (Cat # AM4300, 1:5,000, Abcam) in 1X TBST containing 5% BSA and with primary antibodies against total Akt (Cat # 9272S, 1:1,000, Cell Signaling), total GSK3α/β (Cat # 5676S, 1:1,000, Cell Signaling) or Sox2 (Cat # 14962S, Cell Signaling 1:1,000) in 1X TBST containing 5% BSA for phosphoproteins and 1X TBST containing 5% milk for the remaining proteins, overnight at 4°C with gentle rocking. Membranes were then washed three times for 5 minutes each with 1X TBST and incubated with secondary antibodies, 1:2000 anti-mouse or anti-rabbit horseradish peroxidase (HRP; Cell Signaling) in 5% milk TBST for two hours at room temperature with gentle rocking. Membranes were washed again three times for 5 minutes each with 1X TBST. Pierce ECL Plus Western Blotting Substrate (Thermo Fisher Scientific) was added to each membrane and incubated at room temperature for 5 minutes. The blots were then used to expose X-ray films (Agfa X-Ray film, VWR, Cat # 11299-020) in a dark room. The bands that developed were scanned and their density was measured using ImageJ software. The ratio of the gene of interest over its endogenous control was calculated and expressed as relative intensity.

To analyze for changes in the expression levels of Yamanaka factors in HK-374 cells five days after exposing them to 8 Gy irradiation, Western blotting was performed using proteins extracted from FACS sorted ZsGreen-negative and ZsGreen-positive cells treated with 8 Gy. The membranes were blotted against Sox2, Oct 4 (Cell Signaling, Cat # 2750S), Klf4 (Cell Signaling, Cat # 4038S), cMyc (Cell Signaling, Cat # 5605S) and Nanog (Cell Signaling, Cat # 4903S). GAPDH was used as the loading control.

*Immunohistochemistry*

Brains were explanted, fixed in formalin for twenty-four hours and embedded in paraffin. 4 μm thin sections were baked for one hour in an oven at 65° C, de-waxed in 2 successive Xylene baths of 5 minutes then hydrated for 5 minutes using an alcohol gradient (ethanol 100%, 90%, 70%, 50%, 25%). The slides were incubated in 3% hydrogen peroxide/methanol solution for 10 minutes. Antigen retrieval was performed using Heat Induced Epitope Retrieval in a citrate buffer (10 mM sodium citrate, 0.05% tween20, pH 6) with heating to 95°C in a steamer for 25 minutes. After cooling down, the slides were incubated with the primary antibody against Ki67 (Abcam Cat #15580, 1:200) or β-catenin (Cell Signaling Cat #9562S, 1:200) for 45 minutes at room temperature. The slides were then incubated with Dako EnVision+ System –HRP Labeled Polymer anti-rabbit (Dako, K4003) at room temperature for 30 minutes, rinsed, then incubated with DAB (Betazoid DAB Chromogen kit, Biocare medical) for 10 minutes. Tissues were counterstained with Harris modified Haematoxylin (Fisher scientific, 30 seconds), dehydrated via an alcohol gradient (ethanol 25%, 50%, 70%, 90%, 100%) and soaked twice into Xylene. A drop of Fluoromount aqueous mounting media (Sigma) was added on the top of the section before covering up with a coverslip.

**Supplementary Table 1**

| Gene name | Primer sequence (5’ -3’) |
| --- | --- |
| cMYC | Forward: CACTGTCCAACTTGACCCTCTTG  Reverse: CGTCTCCACACATCAGCACAA |
| KLF4 | Forward: GGTCCGACCTGGAAAATGCT  Reverse: ACCAGGCACTACCGTAAACACA |
| OCT4 | Forward: CATAGTCGCTGCTTGATCGCTTG  Reverse: GAGAACCGAGTGAGAGGCAACC |
| SOX2 | Forward: TTGCGTGAGTGTGGATGGGATTGGTG  Reverse: GGGAAATGGGAGGGGTGCAAAAGAGG |
| NANOG | Forward: TGCGTCACACCATTGCTATTCTTC  Reverse: AATACCTCAGCCTCCAGCAGATG |
| DRD1 | Forward: GGGCTGTTGCTTTTCTGGTG  Reverse: TAGGTTGGGCTTGACGTGAG |
| DRD2 | Forward: TGTACAATACGCGCTACAGCTCCA  Reverse: ATGCACTCGTTCTGGTCTGCGTTA |
| DRD3 | Forward: GTGGTGTCCTTCTACCTGCC  Reverse: GAGAGAGGGTTTGTTGGGGG |
| DRD4 | Forward: ATCCTCACCTGCTCCTCGGTT  Reverse: CCTGGATAGAAGGACGGGCA |
| DRD5 | Forward: CCTCACTCAACCCCGTCATC  Reverse: CAGCTGCGATTTCCTTGTGG |
| DMKN | Forward: CCGACTCTGGGAGGATTTCA  Reverse: TTCTGATCGTCTCTGCCTGC |
| PLA2G3 | Forward: AATCAGCACGACTCCATCTCG  Reverse: CCAGGAGGCATTGTAGAAGGT |
| PCSK9 | Forward: ATGGTCACCGACTTCGAGAAT  Reverse: GTGCCATGACTGTCACACTTG |
| BAI1 | Forward: GCGGCGCTACACTCTCTAC  Reverse: GCACCTCGTCGAAGCTCTC |
| IL36RN | Forward: ACTCGGCATTGAAGGTGCTTT  Reverse: GGACCACGCTGATCTCTT |
| MMP3 | Forward: CTGGACTCCGACACTCTGGA  Reverse: CAGGAAAGGTTCTGAAGTGACC |
| HS3ST2 | Forward: CCAAGCTGATCGTGGTTGTG  Reverse: CCTCAAAGGTCGGGATGTCG |
| FOLR3 | Forward: CGCAAAGAGCGCATTCTGAAC  Reverse: CTGGGCTGAGTCAAACCACA |
| PRUNE2 | Forward: ACCAGTGCTGAACATACCAAG  Reverse: ACTGCTGCCAACAAGTGTTAT |
| FAM111B | Forward: GCCCTTGAAATGCAGAATCCA  Reverse: GCTGTAAACACACTACGGTCTAA |
| SOX7 | Forward: TCGACGCCCTGGATCAACT  Reverse: CTGGGAGACCGGAACATGC |
| PER2 | Forward: CTTCAGCGATGCCAAGTTTGT  Reverse: CGGATTTCATTCTCGTGGCTTT |
| ORC1 | Forward: ACCGAGATTCACATCCAGATTGG  Reverse: CGAGCACGTTTCTTAGGAGGA |
| ADAMTS15 | Forward: TGCGACGCTGCTTCTATTCTG  Reverse: CCTCGGTAGCCAAAGGCTC |
| RRM2 | Forward: CACGGAGCCGAAAACTAAAGC  Reverse: TCTGCCTTCTTATACATCTGCCA |
| DTL | Forward: TAAAAGCTGGTGAGCTGATTGG  Reverse: TCTTCCACCCGTACAGAATACA |
| WDR76 | Forward: AGCTACAACCCAAGAGAACGG  Reverse: CCCGAAAAATCCAGGGATGGT |
| MCM10 | Forward: CCCCTACAGACGATTTCTCGG  Reverse: CAGATGGGTTGAGTCGTTTCC |
| LINC00707 | Forward: TCACCTTCGGCCCATTTCTC  Reverse: TGGTGAATAGTGGAGGGGGT |
| IPO8 | Forward: CGAAGTTGCGGATTGCAG  Reverse: GAATTCCACATGGTCAGAGACT |
| TBP | Forward: TGCACAGGAGCCAAGAGTGAA  Reverse: CACATCACAGCTCCCCACCA |
| PPIA | Forward: ATGCTGGACCCAACACAAAT  Reverse: TCTTTCACTTTGCCAAACACC |

**Supplementary Figure 1**

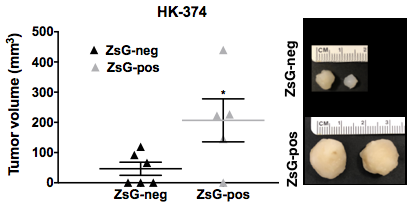

Sorted ZsGreen-cODC-negative and ZsGreen-cODC-positive HK-374 cells were injected subcutaneously in the NSG mice at a fixed number of 5000 cells per 100 uL mixed with matrigel at a ratio of 1:1. The study was terminated when the tumor volume in one of the experimental mice reached the protocol end point. The tumors from both groups were excised and their measurements were used to calculate the tumor volumes.

**Supplementary Figure 2**

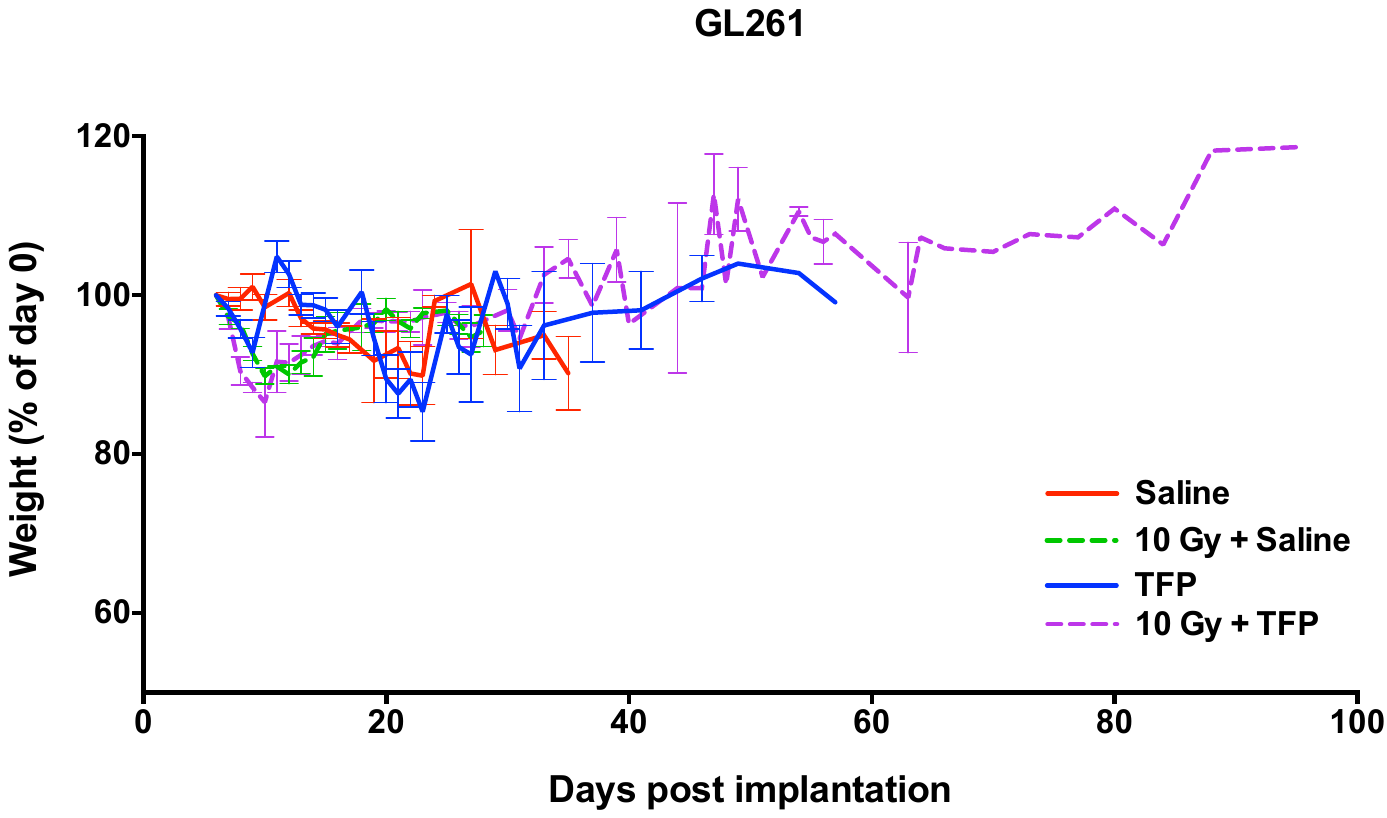

**A**

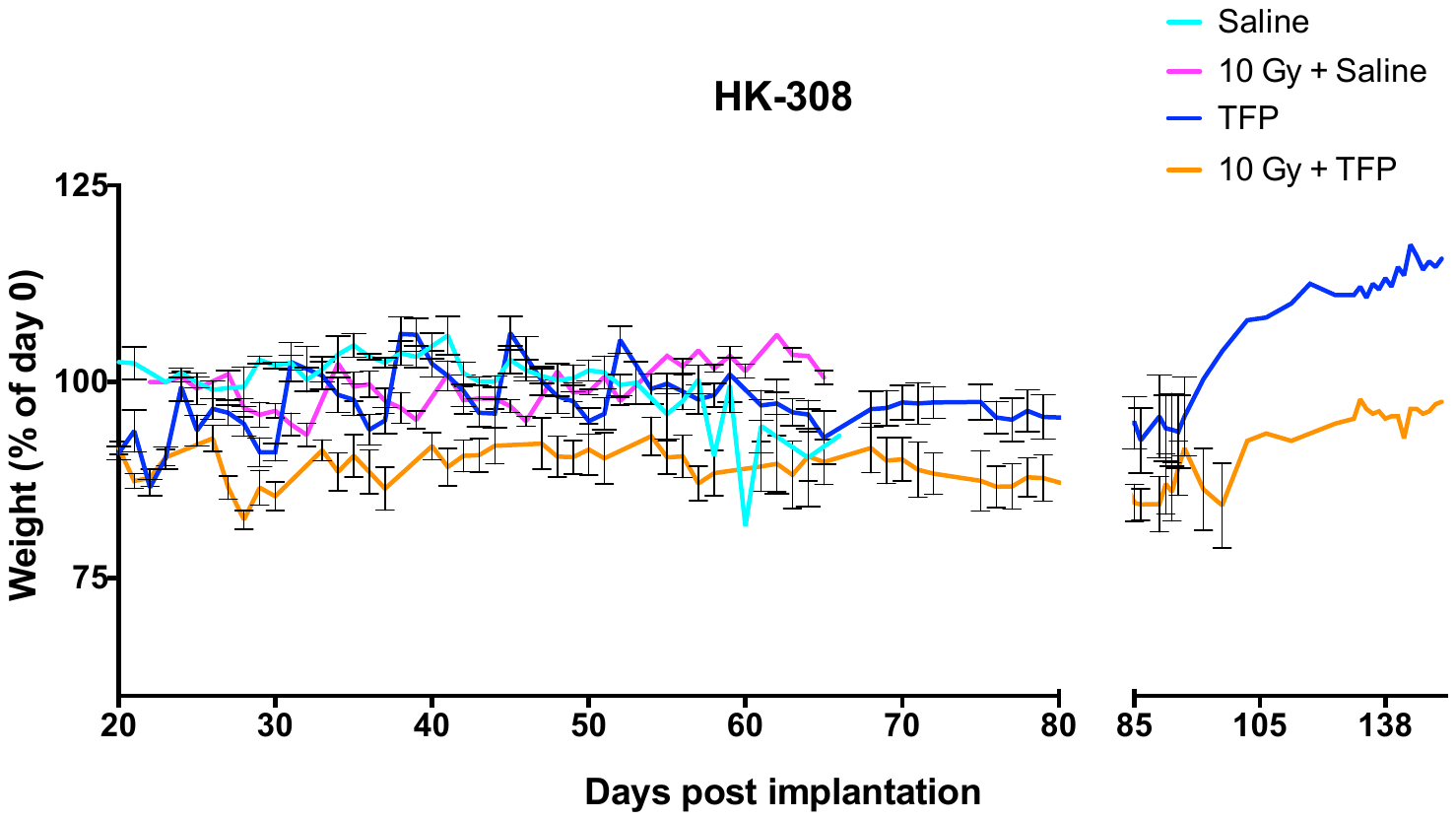

**C**

**B**

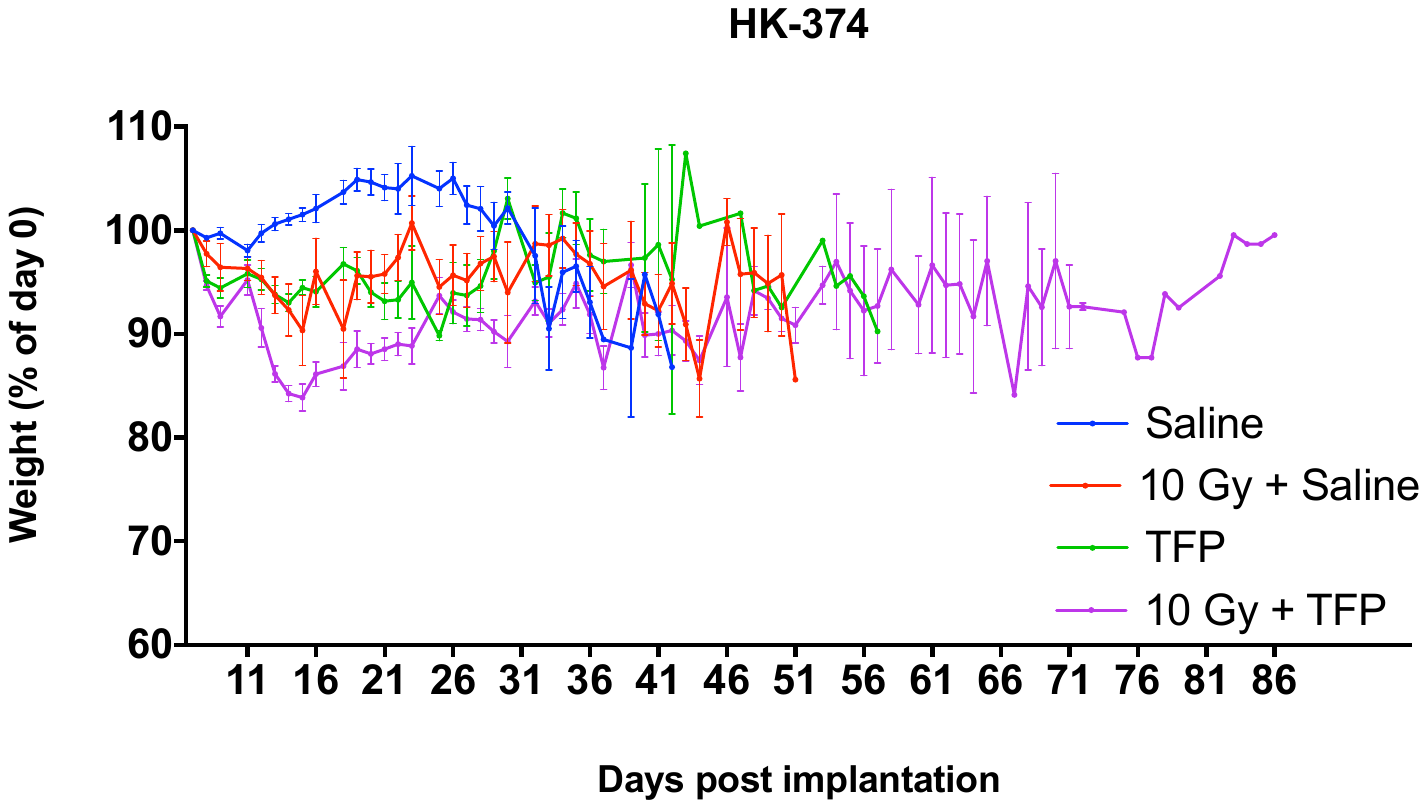

6–8-week-old C57BL/6 mice, or NOD-*scid* IL2Rgamma^null^ (NSG) originally obtained from The Jackson Laboratories (Bar Harbor, ME) were re-derived, bred and maintained in a pathogen-free environment in the American Association of Laboratory Animal Care-accredited Animal Facilities of Department of Radiation Oncology, University of California (Los Angeles, CA) in accordance to all local and national guidelines for the care of animals. 2x10^5^ GL261-Luc and 3x10^5^ HK-308-Luc or HK-374-Luc cells were implanted into the right striatum of the brains of mice using a stereotactic frame (Kopf Instruments, Tujunga, CA) and a nano-injector pump (Stoelting, Wood Dale, IL). Injection coordinates were 0.5mm anterior and 2.25mm lateral to the bregma, at a depth of 2.8mm from the surface of the brain. Tumors were grown for 3 (HK-374), 7 (GL261) or 21 (HK-308) days after which successful grafting was confirmed by bioluminescence imaging. Mice that developed neurological deficits requiring euthanasia were sacrificed. Mice-bearing tumors were injected intra-peritoneally on a 5-days on / 2-days off schedule for 3 weeks (GL261), and continuously (HK-374 and HK-308) either with TFP or saline. TFP was dissolved in sterile saline at a concentration of 2.5 mg/mL. All animals were treated with 20 mg/kg TFP. Weights of the animals were recorded every day.
